# Chemoproteomics of microbiota metabolites reveals small-molecule agonists for orphan receptor GPRC5A

**DOI:** 10.1101/2021.12.16.472979

**Authors:** Xiaohui Zhao, Kathryn R. Stein, Victor Chen, Matthew E. Griffin, Howard C. Hang

## Abstract

The microbiota generates diverse metabolites that can engage multiple pathways to modulate host physiology and disease, but their protein targets and mechanism(s) of action have not been fully elucidated. To address this challenge, we focused on indole-3-acetic acid (IAA), a prominent microbiota metabolite, and developed IAA-based chemical reporters for proteomic studies. We discovered that IAA interacts with many proteins in host cells, including small-molecule transporters, receptors and metabolic enzymes. Notably, our functional studies revealed that IAA binds to orphan G protein-coupled receptors such as GPRC5A, but only aromatic monoamines were capable of inducing GPRC5A signaling. Functional profiling of microbiota uncovered specific bacterial species and enzymes that generate GPRC5A agonists. Finally, biochemical characterization of GPRC5A activation identified more potent synthetic agonists as well as key amino acid residues involved in ligand binding. These studies highlight the utility of chemoproteomics to dissect protein targets and mechanisms of action for microbiota metabolites.

## INTRODUCTION

The human microbiota directly modulate host immunity and generate diverse metabolites that engage multiple pathways to regulate host physiology.^1–3^ For example, microbe-associated molecular patterns (MAMPs) such as lipopolysaccharides (LPS) can engage toll-like receptors (TLRs) to maintain intestinal epithelial homeostasis^4^ and modulate infant autoimmunity^5^. In addition, microbiota-derived cyclic dinucleotides^6^ and peptidoglycan fragments^7,8^ initiate host immune response and enhance the efficacy of immune checkpoint inhibitor for cancer therapy^9,10^. Beyond generating MAMPs that engage canonical pattern recognition receptors (PRRs), microbiota metabolism of dietary components (e.g., polysaccharides, amino acids and lipids)^11^, host metabolites (e.g. bile acids)^12^ and therapeutic drugs^13^ can produce diverse molecules in micromolar to millimolar levels, which have local and systemic effects on host health and disease. Notably, aromatic amino acids (aAAs) are transformed by microbiota to diverse metabolites, such as aromatic hydrocarbons, acids and amines.^14,15^ These microbiota-derived aromatic compounds have been suggested to engage different classes of protein targets within host cells, including nuclear receptors^16,17^ and G protein-coupled receptors (GPCRs)^18,19^, but their other potential protein targets and mechanisms of action have not been fully elucidated.

Advances in chemical proteomics provide an excellent opportunity to characterize protein targets of microbiota metabolites and elucidate their mechanisms of action in host pathways.^20–22^ The derivatization of microbiota metabolites with bioorthogonal tags for detection and affinity enrichment as well as photoaffinity moieties for crosslinking enables identification of both covalently and non-covalently labeled proteins for subsequent functional studies.^23,24^ To showcase the utility of chemoproteomics for microbiota metabolite studies, we developed metabolic (alk-IAA) and photoaffinity (x-alk-IAA) reporters of indole-3-acetic acid (IAA), a prominent microbiota-generated metabolite^25,26^ that exhibits significant effects on host physiology through engaging multiple pathways.^27,28^ From chemoproteomic profiling of IAA-interacting proteins in intestinal epithelial cells, we identified many candidate protein targets including smallmolecule transporters, immune sensors and GPCRs, highlighting the utility of chemoproteomics to reveal potentially novel mechanisms of action for microbiota metabolites.

Our functional studies revealed that the microbiota generate small-molecule aromatic monoamine agonists for orphan receptor GPRC5A. Specific microbiota species and enzymes, such as *Enterococcus* species that encode tyrosine decarboxylases, are capable of activating GPRC5A. In addition, our chemical and biochemical characterization of GPRC5A activation yielded potent synthetic agonists such as 7-fluorotryptamine (7FTA) as well as key amino acid residues for ligand binding. Given the orphan status of GPRC5A and its association with cancer^29–31^ and immunity^32^, these studies showcase how chemoproteomic analysis of microbiota metabolites enables the identification of unpredicted protein targets such as orphan GPCRs and reveals novel mechanism(s) of action between specific microbiota species and metabolic enzymes, microbiota metabolites, and host pathways for potential therapeutic development.

## RESULTS

### Chemoproteomics reveals IAA-interacting proteins

To investigate the protein targets of microbiota metabolites, we initially focused on IAA and generated a metabolic reporter functionalized with an alkyne chemical tag (alk-IAA) as well as a photoaffinity reporter that also contains a diazirine crosslinker (x-alk-IAA) (**Fig. 1a**). To examine whether these chemical reporters function as active analogs of IAA, we tested them on aryl hydrocarbon receptor (AhR), a previously described IAA protein target,^16,33^ using the xenobiotic response element (XRE)-driven luciferase reporter assay in HEK293T cells. Both alk-IAA and x-alk-IAA were able to activate AhR, although less efficient than IAA and a potent ligand 2,3,7,8-tetrachlorodibenzo-p-dioxin (TCDD) (**Fig. 1b**). UV irradiation of the chemical reporter treated cells revealed that x-alk-IAA, but not alk-IAA, can photocrosslink HA-tagged AhR, which was immunoprecipitated, reacted with azide-rhodamine and analyzed by in-gel fluorescence profiling (**Fig. 1c**). Cells treated with DMSO, alk-IAA or x-alk-IAA without UV irradiation did not label HA-AhR. Taken together, these experiments indicate that our IAA chemical reporters function as reasonable analogs of IAA in mammalian cells.

**Fig. 1.**
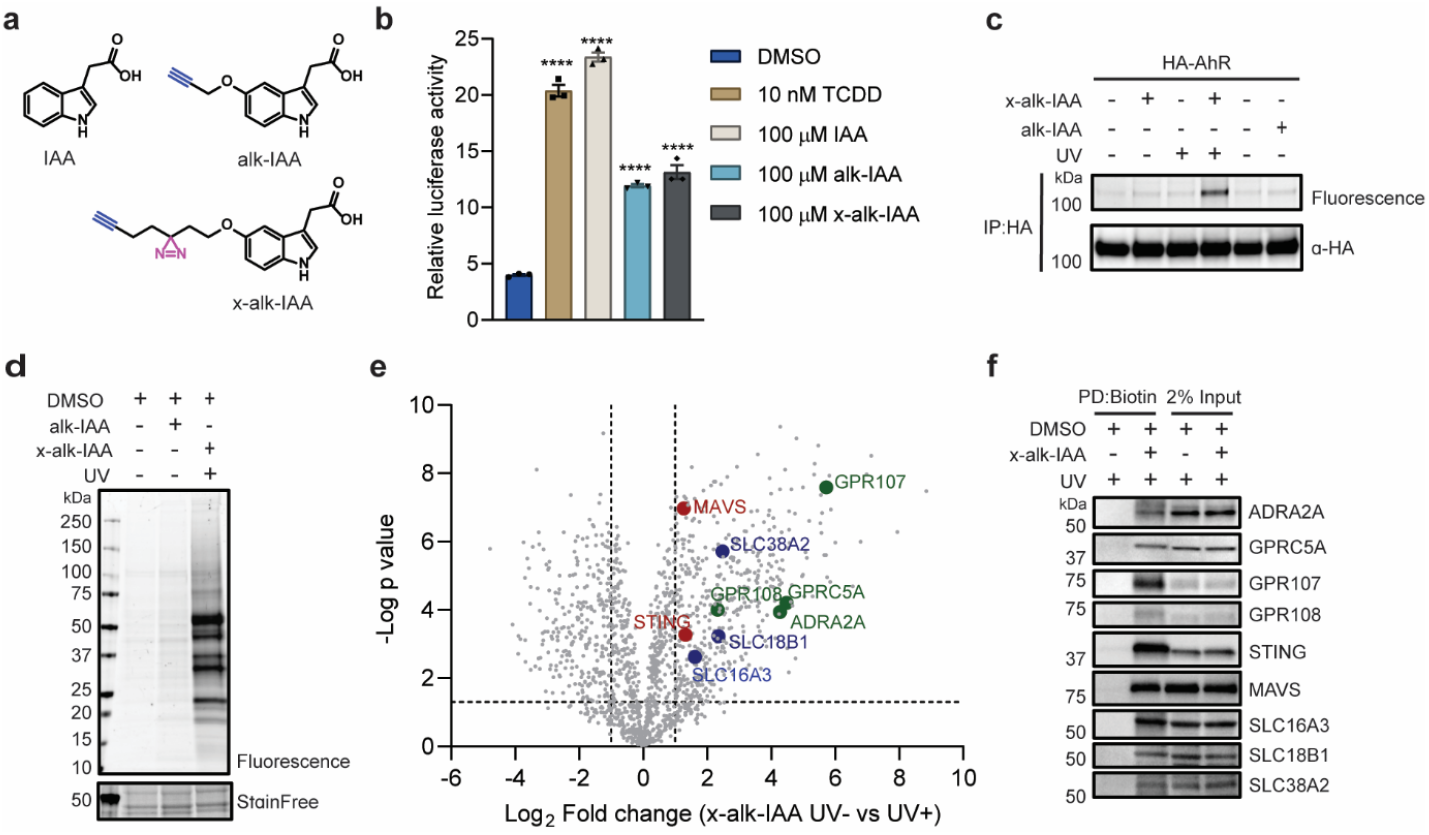
Chemoproteomic analysis of indole-3-acetic acid-interacting proteins in mammalian cells. **a**, Chemical reporters based on indole-3-acetic acid (IAA) for metabolic labeling (alk-IAA) and photoaffinity profiling (x-alk-IAA) of IAA-interacting proteins. **b**, Activity of TCDD, IAA, alk-IAA and x-alk-IAA on aryl hydrocarbon receptor (AhR) and xenobiotic response element (XRE)-luciferase reporter in HEK293T cells. **c**, Photoaffinity crosslinking of x-alk-IAA (100 μM) to HA-tagged AhR in HEK293T cells reveled by rhodamine labeling. **d**, In-gel fluorescent profiling of IAA-interacting proteins in HT-29 cells using 0.1% DMSO, 100 μM alk-IAA or 100 μM x-alk-IAA. **e**, Volcano plot of the identified proteins from HT-29 cells by chemical proteomics using x-alk-IAA (100 μM) and UV irradiation in comparison with samples without UV. p value is 0.05, two folds change in four replicates. Key hits are highlighted in GPCRs (green), smallmolecule transporters (blue) and immunity-associated proteins (red). **f,** Western blot analysis of x-alk-IAA (100 μM) photo-crosslinking proteins. Input indicates 2% of total HT-29 cell lysates. Data indicates mean with SEM from three replicates. One-way ANOVA using Dunnett’s multiple comparisons test. **** indicates p-value < 0.0001.

We next applied the IAA-based chemical reporters to HT-29 cells, a human colorectal adenocarcinoma epithelial cell line, to identify potential microbiota metabolite protein targets using chemoproteomics (**Supplementary Fig. 1a**). The alk-IAA reporter may be metabolically processed in cells and covalently incorporated onto proteins, while the photoaffinity reporter x-alk-IAA enables the photocrosslinking of non-covalent protein targets with UV. HT-29 cells were thus incubated with these IAA chemical reporters with or without UV treatment and total cell lysates were reacted with azide-modified reagents for in-gel fluorescence profiling or label-free quantitative proteomics. HT-29 cells treated with alk-IAA revealed a few specifically labeled proteins compared to DMSO treated samples (**Fig. 1d**). In contrast, we observed many labeled proteins between 20 to 75 kDa with x-alk-IAA and UV irradiation compared to controls (**Fig. 1d**). Labeling of x-alk-IAA proteins correlated with increasing concentration of x-alk-IAA (**Supplementary Fig. 1b**). We therefore reacted the cell lysates with azide-biotin, performed neutravidin affinity enrichment and analyzed the recovered proteins by label-free quantitative proteomics. A total number of 413 proteins were detected in x-alk-IAA and UV-treated samples in comparison to the without UV control (p value < 0.05, two folds change in difference), visualized by volcano plot (**Fig. 1e, Supplementary Table 1**). The majority of these proteins were additionally found when comparing x-alk-IAA and UV-treated samples versus a DMSO control (**Supplementary Fig. 1c,d, Supplementary Table 1**). Notably, the 18 significant proteins recovered from alk-IAA metabolic labeling were also identified with the x-alk-IAA (**Supplementary Fig. 1d, Supplementary Table 1**). Gene ontology (GO) analysis of the hits revealed many cytosolic metabolic enzymes as well as membrane proteins that localize on organelle membranes including the endoplasmic reticulum (ER), mitochondria, Golgi apparatus and lysosome (**Supplementary Fig. 1e**). KEGG pathway analysis of these hits revealed proteins associated with steroid and *N*-glycan biosynthesis, valine, leucine and isoleucine degradation, lipid (e.g., glycerolipid, glycerophospholipid, sphingolipid, cholesterol) and amino acids (e.g., tryptophan and histidine) metabolism (**Supplementary Fig. 1f**). In addition to these metabolic enzymes, we identified proteins involved in different functional classes that include small-molecule transporters (SLC16A3, SLC18B1 and SLC38A2), innate immunity effectors (STING and MAVS) and GPCRs (ADRA2A, GPRC5A, GPR107 and GPR108) (**Fig. 1g**). The x-alk-IAA photocrosslinking of these proteins were validated by western bot analysis using specific antibodies (**Fig. 1h**). The recovery of STING with x-alk-IAA is consistent with prior reports, which demonstrate that this innate immune sensor can interact with aromatic compounds containing a free acid moiety.^34,35^ Conversely, x-alk-IAA photocrosslinking to GPRC5A, GPR107 and GPR108 was striking, as no small molecule ligands were previously reported for these orphan or putative GPCRs, which highlights the utility of chemoproteomics to identify unpredicted protein targets.

The GPCRs crosslinked by x-alk-IAA are interesting candidate microbiota metabolite protein targets. Alpha-2A adrenergic receptor (ADRA2A) responds to the endogenous ligands including epinephrine and norepinephrine to transduce cellular signaling in the sympathetic nervous system^36^ and has been suggested to sense the microbiota-derived metabolite, phenylacetylglutamine^37^. In contrast, GPR107 and GPR108 consist of a 7-transmembrane helix and mainly localize on the *trans*-Golgi network,^38,39^ but no ligands have been reported and functions as GPCRs are unclear.^40^ GPRC5A is a retinoic acid-induced protein [also termed retinoic acid-induced gene 3 (RAI3) or retinoic acid-induced gene 1 (RAIG1)] that is highly expressed in the lung and gastrointestinal tract and associated with inflammation and cancer progression.^41^ GPRC5A mainly localizes to the plasma membrane and endoplasmic reticulum (ER) and is categorized as a class C orphan GPCR. GPRC5A has a short N-terminus that is atypical of class C GPCRs, which usually contain long extracellular N-termini as a ligand-binding domain and a cysteine-rich domain^42^. As GPRC5A is implicated as an immune-associated tumor suppressor with no reported ligands, we investigated whether specific microbiota species and metabolites may function as agonists of this orphan GPCR.

### Aromatic monoamines activate GPRC5A signaling

To explore whether IAA and indole-derived metabolites modulate GPCR activity of the identified proteins, we utilized PRESTO-Tango^43^, a cell-based reporter assay that relies on β-arrestin 2 recruitment and transcriptional activation. We tested IAA and a collection of indole molecules, including indole (Id), tryptophan (Trp), indole propionic acid (IPA), indole acrylic acid (IAcA), tryptamine (TA), indole aldehyde (IAld), indole-3-carbinol (I3C), and x-alk-IAA photoaffinity reporter (XAI) (**Fig. 2a**). For ADRA2A, TA exhibits weak effects in comparison with its endogenous ligand epinephrine (Epi) and synthetic agonist clonidine (Clo) (**Fig. 2b**). Notably, amongst the collection of indole molecules screened, TA activated GPRC5A most effectively (**Fig. 2c**) with an EC_50_ of 88.9 μM (**Fig. 2d**). In addition, TA competes with x-alk-IAA photocrosslinking to GPRC5A and ADRA2A (**Supplementary Fig. 2a,b**). Interestingly, IAA also competed with x-alk-IAA photocrosslinking (**Supplementary Fig. 2a,b**) but did not activate GPRC5A (**Fig. 2c,d**). These results suggest that IAA may bind to GPRC5A with low-affinity, but is incapable of inducing downstream signaling compared to TA. To further investigate TA interaction with proteins, we synthesized a TA photoaffinity reporter x-alk-TA (**Fig. 2e**) and found that it more effectively photocrosslinked endogenous GPRC5A in HT-29 cells (**Fig. 2f**) and FLAG-tagged GPRC5A expressed in HEK293T cells in comparison to x-alk-IAA (**Fig. 2g**), suggesting TA may bind GPRC5A stronger than IAA. These data suggest that additional microbiota aromatic monoamines may also activate GPRC5A signaling. We therefore evaluated aromatic monoamines such as phenethylamine (PEA), tyramine and histamine. PEA showed comparable activity to TA at 100 μM, whereas tyramine and histamine only exhibited minimal or no activity on GPRC5A signaling (**Fig. 2h**). These results demonstrate that aromatic metabolites bind to GPRC5A, but only aromatic amines can effectively induce signaling, which is the first indication for smallmolecule agonists of GPRC5A and deorphanizes this class C GPCR.

**Fig. 2.**
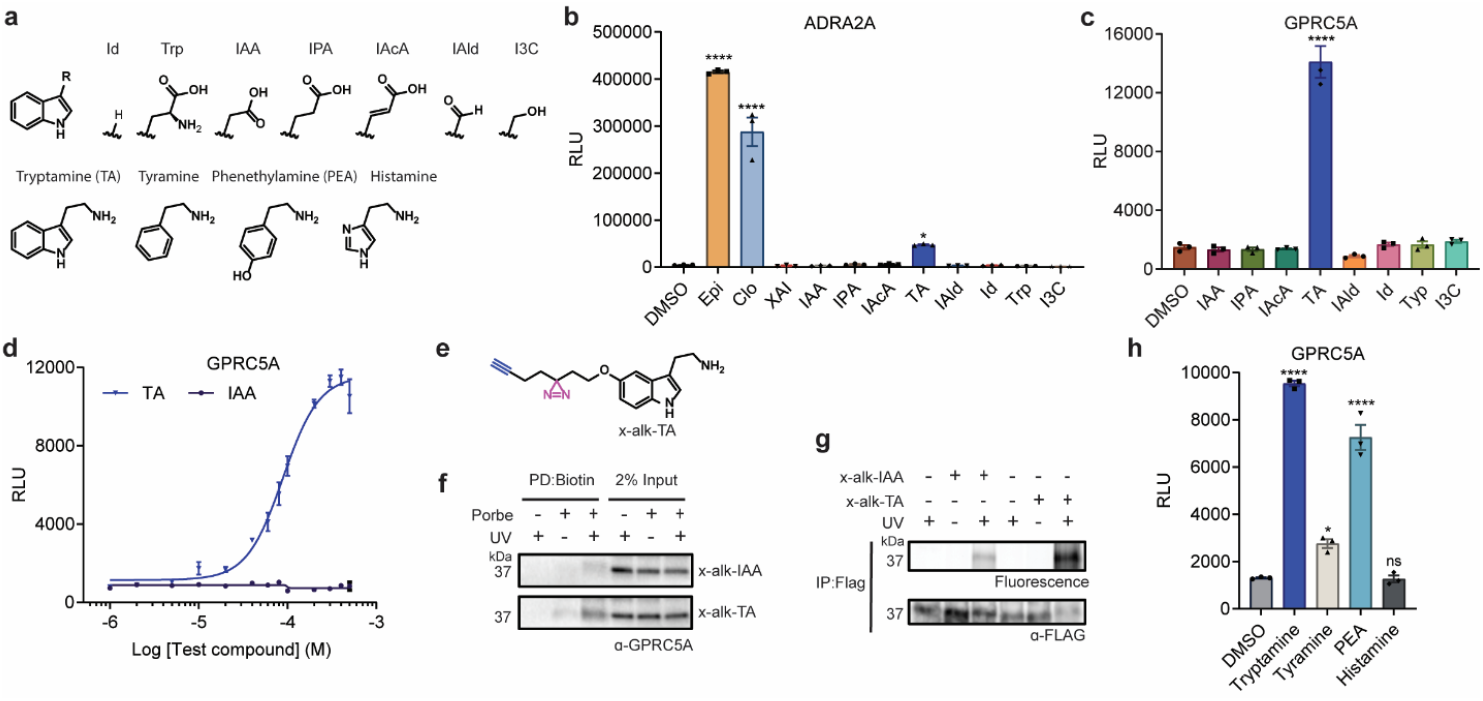
Aromatic monoamines activate GPRC5A signaling. **a**, Chemical structures of indole derivatives and aromatic monoamines. **b,c**, PRESTO-Tango assay for screening a collection of indole derivatives (100 μM) on ADRA2A (b) and GPRC5A (c) activation. 0.1% DMSO, 1 μM epinephrine (Epi), 1 μM clonidine were used as controls. **d**, Dose-response curves of GPRC5A activation upon IAA and TA treatment. **e**, Chemical structure of TA photoaffinity reporter x-alk-TA. **f,** Pull-down of endogenous GPRC5A from HT-29 cells that were photo-crosslinked by x-alk-IAA or x-alk-TA followed by biotin labeling and neutravidin enrichment. **g**, Photoaffinity crosslinking of x-alk-IAA or x-alk-TA to FLAG-tagged GPRC5A in HEK293T cells identified with rhodamine labeling. **h**, PRESTO-Tango assay for examine aromatic monoamines (100 μM) in GPRC5A activation. Data indicates mean with SEM from three replicates. One-way ANOVA using Dunnett’s multiple comparisons test. ****p<0.0001, *p<0.05, ns indicates not significant.

### Microbiota species containing aAAs decarboxylases activate GPRC5A signaling

We next explored GPRC5A activating microbiota species and enzymes that could generate aromatic monoamines. TA, PEA and tyramine are aromatic monoamines produced by the metabolic processing of aAAs by bacteria and human. For example, two common Firmicutes, *Ruminococcus gnavus* and *Clostridium sporogenes* converts Trp to TA through reductive metabolism by the Trp decarboxylase (TrpDC).^44^ Alternatively, PEA is known as a product of bacterial species including *Enterococcus faecalis*^45^ and *Morganella morganii*^9^, whereas tyramine has been identified in the growth media of different *Enterococcus* and *Lactobacillus* species.^45,46^ To identify microbiota species that can activate GPRC5A, we examined the growth media of different bacterial species and strains predicted to metabolize aAAs into their corresponding monoamines. The bacterial growth media were subjected to 3 kDa MW-cutoff membrane filtration to remove secreted proteins and retain low molecular weight metabolites. Amongst the bacterial species we initially screened, 3 kDa MW-cutoff membrane filtered growth media from *R. gnavus, M. morganii, E. faecium* and *E. faecalis* effectively activated GPRC5A (**Fig. 3a**). However, the membrane filtered growth media from *C. sporogenes* and *E. coli* exhibited no GPRC5A activation beyond their control growth media alone (**Fig. 3a**). To determine whether these bacterial species generate aromatic monoamines, we performed targeted LC-MS analysis of the 3 kDa MW-cutoff membrane filtered growth media (**Fig. 3b**). Indeed, *R. gnavus, E. faecium* and *E. faecalis* generate high micromolar levels of GPRC5A aromatic monoamine agonists (**Fig. 3b**). Instead of generating different aromatic monoamines, *M. morganii* primarily produced PEAs at higher levels ~500 μM (**Fig. 3b**). *C. sporogenes* also generated aromatic monoamines, but at lower micromolar levels (**Fig. 3b**), which may not be sufficient to activate GPRC5A signaling in the PRESTO-Tango assay (**Fig. 3a**). *E. coli* did not produce significant levels of aromatic monoamines that were detectable by our fractionation and LC-MS conditions (**Fig. 3b**). Based on the *E. faecium* and *E. faecalis* results and the importance of *Enterococcus*-host interactions, we evaluated additional *Enterococcus* strains and species, which revealed that *E. faecium* and *E. faecalis* strains as well as *E. durans, E. hirae*, and *E. mundtii* membrane filtered growth media activated GPRC5A (**Fig. 3c**). However, the growth media from *E. gallinarum* exhibited no significant effect on GPRC5A activation (**Fig. 3c**).

**Fig. 3.**
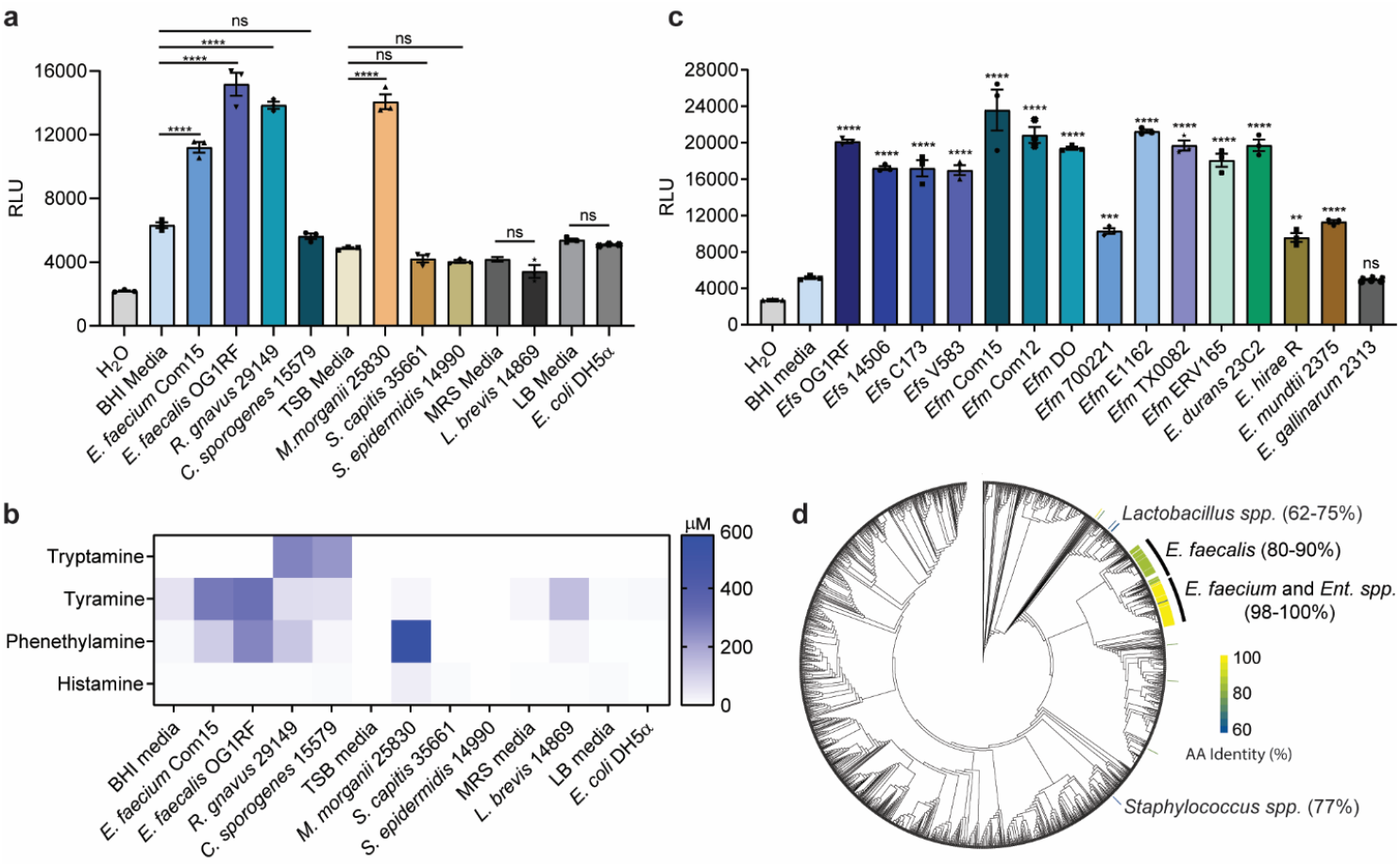
Specific microbiota species and strains produce aromatic monoamines that activate GPRC5A signaling. **a**, GPRC5A activation by PRESTO-Tango analysis of bacterial cultures of representative human microbiota species. **b**, Heat map of aromatic monoamines concentration in different bacterial cultures determined by LC-MS analysis. **c**, GPRC5A activation by PRESTO-Tango analysis of different *Enterococcus* species and strains. **d**, Cladogram of Human Microbiome Project isolates organized by 16S rRNA homology with heatmap indicating amino acid (AA) sequence identity of putative tyrosine decarboxylase orthologs in *Enterococcus, Lactobacillus*, and *Staphylococcus* species. *Ent*. spp. indicates the *Enterococcus* isolates without a specific species annotation. Data indicates mean with SEM from three replicates. One-way ANOVA using Tukey’s multiple comparisons test in comparison with respective media (a) or Dunnett’s multiple comparisons test in comparison with BHI media (c). ****p<0.0001, ***p<0.001, **p<0.01 *p<0.05, ns indicates not significant.

To further investigate GPRC5A activating microbiota species, we performed a bioinformatic searching and phylogenetic analysis of bacterial amino acid decarboxylases. While TrpDC activity has been reported for *R. gnavus*^44^, no clear orthologs greater than 60% protein sequence similarity of this enzyme are present in *Enterococcus*. However, *E. faecium* contains a predicted Tyr decarboxylase (TyrDC) (**Supplementary Fig. 3**) that is highly conserved in the strains of *E. faecium, E. faecalis*, and other *Enterococcus* species identified and sequenced from the Human Microbiome Project (HMP)^47^ **(Fig. 3d, Supplementary Table 2**). *E. gallinarum* strain does not appear to have a TyrDC ortholog, but contains a Gln decarboxylase (**Supplementary Fig. 3**) that may explain its inability to activate GPRC5A (**Fig. 3c**). Other microbiota species such as *Staphylococcus* and *Lactobacillus* encode proteins with more than 60% protein similarity to *E. faecium* TyrDC. However, the 3 kDa MW-cutoff membrane filtered growth media from *S. capitis*, *S. epidermidis* and *L. brevis* did not induce GPRC5A activation (**Fig. 3c**) and only produced minimal levels of aromatic monoamines or undetectable levels under the conditions we evaluated (**Fig. 3b**). These results indicated that microbiota species which express aAAs decarboxylases and generate high micromolar levels of aromatic monoamines (*R. gnavus*, *M. morganii* and *Enterococcus* spp.) function as GPRC5A agonists. Different bacterial species that contain decarboxylases may exhibit lower activity, distinct specificity and/or require unique conditions to generate sufficient levels of aromatic monoamines to activate GPRC5A.

### Bacterial aromatic amino acid decarboxylases generate GPRC5A agonists

To determine the specific aAAs decarboxylases that are involved in the production of aromatic monoamines and activation of GPRC5A, we performed gain- and loss-of-expression in bacteria as well as *in vitro* activity studies. For these studies, we expressed the *E. faecium* tyrosine decarboxylase (Efm_TyrDC), *R. gnavus* tryptophan decarboxylase (Rgs_TrpDC) and *M. morganii* glutamate or tyrosine decarboxylase (Mmi_Gln/TyrDC) in *E. coli* DH5α (**Supplementary Fig. 4a,b**), examined the 3 kDa MW-cutoff membrane filtered growth media for aromatic monoamines by LC-MS (**Supplementary Fig. 4c-n**) and determined the activation of GPRCA by the PRESTO-Tango assay (**Fig. 4a**). We found that all three enzymes did not generate histamine, but effectively decarboxylated Phe to PEA (**Fig. 4a, Supplementary Fig. 4d,h,l**) and activated GPRCA signaling (**Fig. 4b**). As previously described,^44^ Rgs_TrpDC was able to catalyze the decarboxylation of Trp to generate TA (**Fig. 4a, Supplementary Fig. 4i**) and in our assays activate GPRCA signaling (**Fig. 4b**). Mmi_Gln/TyrDC on the other hand was specific for Phe (**Fig. 4a, Supplementary Fig. 4l**). Efm_TyrDC generated high levels of tyramine (**Fig. 4a, Supplementary Fig. 4a**), but has only a mild effect on GPRC5A signaling (**Fig. 4b**), consistent with our data with tyramine alone (**Fig. 2e**). Efm_TyrDC also generated low levels of TA from Trp (**Fig. 4a, Supplementary Fig. 4e**), which displayed modest activation of GPRC5A (**Fig. 4b**), suggesting this decarboxylase is selective for Tyr and Phe over Trp. Indeed, our analysis of recombinant and purified His6-tagged Efm_TyrDC (**Supplementary Fig. 5a**) demonstrated that this *Enterococcus* aAAs decarboxylase preferentially utilizes Tyr over Phe, but not Trp, as a substrate under conditions we tested (**Supplementary Fig. 5b,c**). Importantly, the reaction products from Efm_TyrDC *in vitro* decarboxylation of Phe and Tyr also activated GPRC5A (**Supplementary Fig. 5d**). Kinetic analysis of Efm_TyrDC activity revealed a *V*_max_ of 0.7483 μM/s and a *K*_M_ value of 280.7 μM with Tyr and *V*_max_ value of 0.04 mM/min and *K*_M_ of 8.5 mM with Phe (**Supplementary Fig. 5e**). These results demonstrate bacterial decarboxylases from diverse microbiota species generate aromatic amines that activate GPRC5A signaling, confirming previous enzymatic activity for Rgs_TrpDC and predicted activity of Mmi_Gln/TyrDC and Efm_TyrDC.

**Fig. 4.**
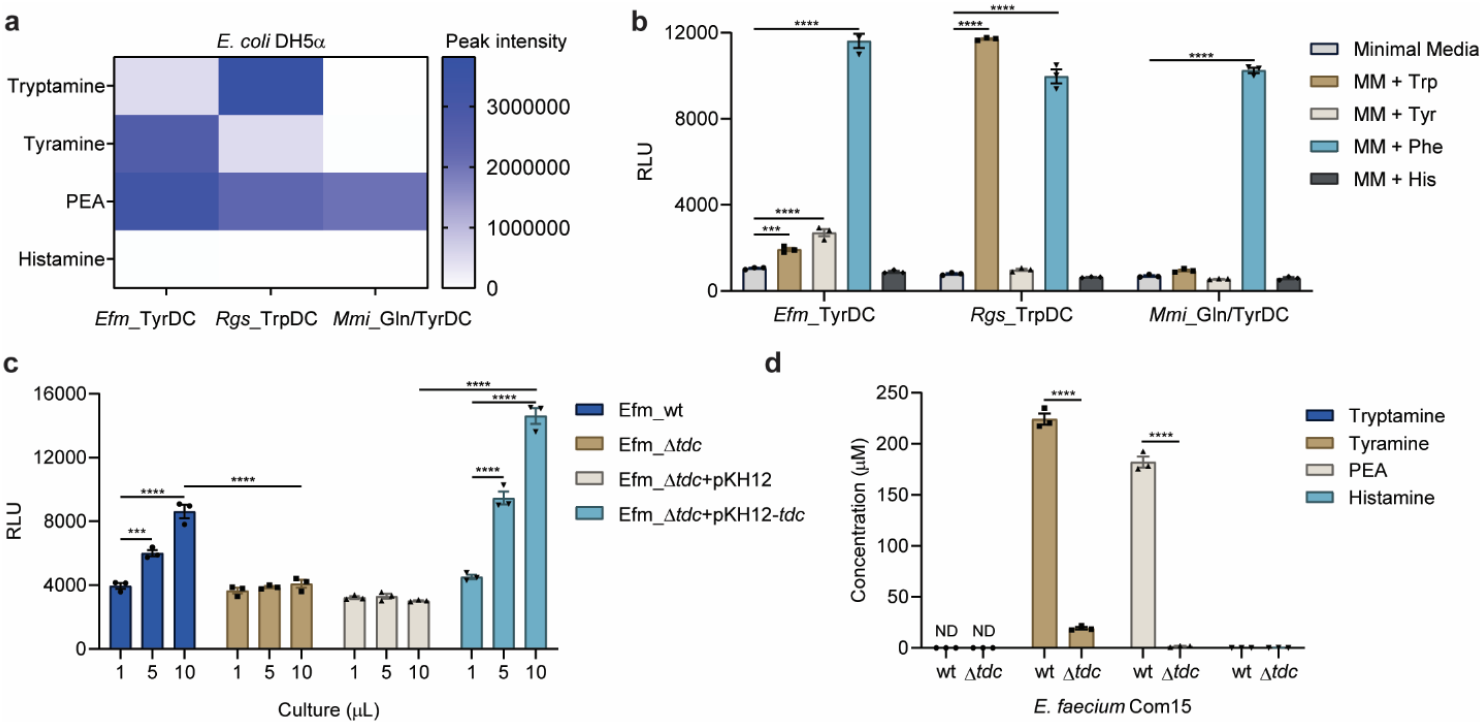
Bacterial decarboxylases generate aromatic monoamines that activate GPRC5A signaling. **a**, LC-MS determination of aromatic monoamines produced by overexpression of *E. faecium* Com15 tyrosine decarboxylase (*Efm*_TyrDC), *R. gnavus* tryptophan decarboxylase (*Rgs*_TrpDC) and *M. morganii* glutamate/tyrosine decarboxylase (*Mmi*_Gln/TyrDC) in *E. coli* DH5α. **b**, PRESTO-Tango assay for comparing bacterial enzyme products on GPRC5A activation. **c**, PRESTO-Tango assay for examining wild-type *E. faecium* Com15 (*Efm*_wt) and the *E. faecium* Com15 *TyrDC* knock-out (*Efm*_Δ*tdc*) cultures with TyrDC (pKH12-*tdc*) complementation or vector control (pKH12) on GPRC5A activation. **d**, LC-MS determination of aromatic monoamines in *Efm_wt* and *Efm*_Δ*tdc* cultures. Data indicates mean with SEM from three replicates. Two-way ANOVA using Tukey’s multiple comparisons test. ****p<0.0001, ***p<0.001, ND indicates not detected.

To determine if Efm_TyrDC is also essential or required to generate aromatic amines in *Enterococcus*, we generated a *E. faecium* Com15 *tyrDC* deletion strain (*Efm_ΔtyrDC*) using recently developed CRISPR-recombineering methods (**Supplementary Fig. 6a,b**)^48^. Deletion of *TyrDC* did not affect *E. faecium* cell growth (**Supplementary Fig. 5c**), but abolished the effect on GPRC5A activation (**Fig. 4c**) as well as tyramine and PEA levels in the growth media (**Fig. 4d**). Complementation of *Efm_ΔtyrDC* strain with TyrDC by overexpression restored and enhanced GPRC5A activation (**Fig. 4c, Supplementary Fig. 6d**). The results demonstrate TyrDC is a key aromatic amino decarboxylase in *E. faecium* Com15, which is highly conserved in many *E. faecium* strains and *Enterococcus* species (**Fig. 3d**) and is capable of generating aromatic monoamines to activate GPRC5A.

### GPRC5A preferentially senses aromatic amines and is more effectively activated by 7-fluorotryptamine

To classify the key determinants of aromatic monoamines and identify more potent GPRC5A agonists, we investigated additional natural and synthetic compounds (**Fig. 5a,b**). We found that the aromatic moiety is essential because ethylamine (EA) was not able to activate GPRC5A (**Fig. 5c**). We next observed decreased and complete loss of the GPRC5A activation with *N*-methyl tryptamine (NMTA) and *N*-acetyl tryptamine (NAcTA), respectively (**Fig. 5c**). These data together with the IAA results indicate that aromatic primary amines are the minimal pharmacophore for GPRC5A activation. We therefore tested additional indole-based primary amines, including indole-3-amine (I3A), indole-3-methylamine (I3mA), α-methyltryptamine (αMTA) and α-ethyltryptamine (αETA) (**Fig. 5a**). Three compounds (i.e., I3mA, αMTA, and αETA) retained activity towards GPRC5A (**Fig. 5c**), demonstrating the tolerance of truncation and branching of the alkyl amine moiety. We then evaluated a collection of tryptamine derivatives bearing various substitutions (methyl, hydroxyl, methoxy, halogen) on distinct positions of the indole ring (**Fig. 5b**). Notably, 7-fluorotryptamine (7FTA) was ~5-fold more effective at inducing GPRC5A signaling than TA at 100 μM (**Fig. 5d**) and more active than other TA derivatives (**Fig. 5d, Supplementary Fig. 7a**), with an EC_50_ value of 7.2 μM (**Fig. 5e**). Concentrations of TA and 7FTA below 100 μM exhibited no toxicity for mammalian cells (**Supplementary Fig. 7b**).

**Fig. 5.**
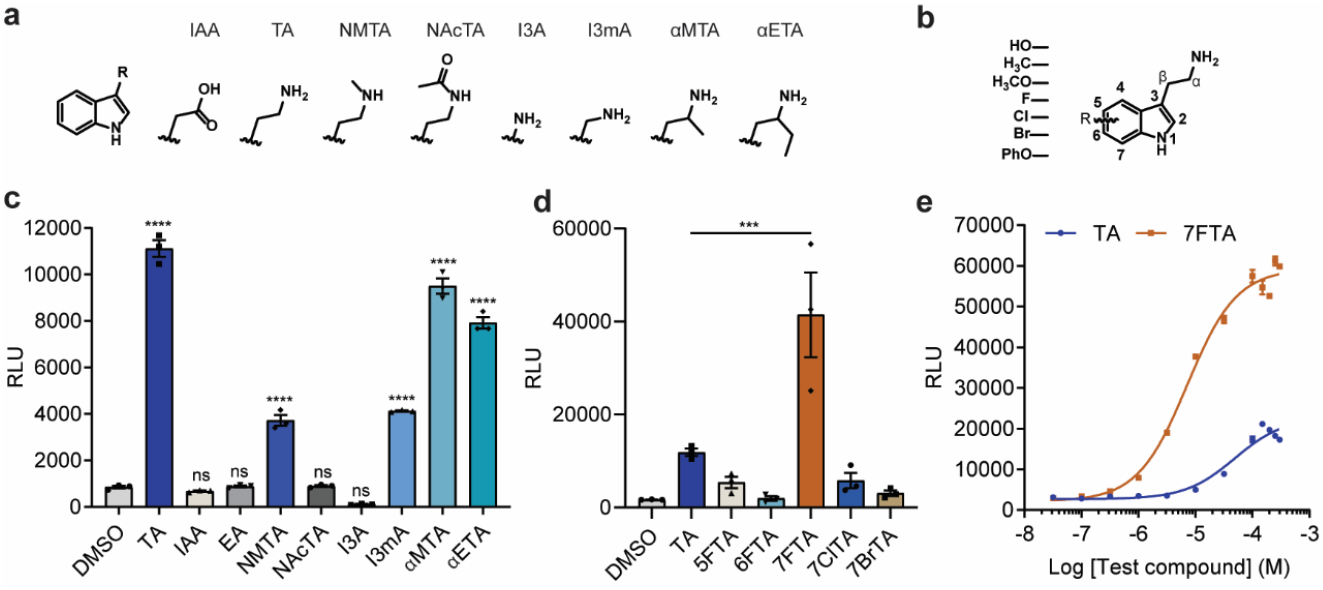
Structure-activity relationship studies of GPRC5A ligands. **a,b**, Chemical structures of indole and tryptamine derivatives. **c-d,** PRESTO-Tango assay for analysis of various indole and tryptamine derivatives (100 μM) on GPRC5A activation. **e**, Dose-response curves of GPRC5A activation upon TA and 7FTA treatment. Data indicates mean with SEM from three replicates. One-way ANOVA using Tukey’s multiple comparisons test. ****p<0.0001, ***p<0.001, ns indicates not significant.

### Aromatic amines bind conserved transmembrane domain of retinoic acid-induced class C orphan GPCRs

To understand how GPRC5A senses aromatic monoamines, we performed site-directed mutagenesis and photocrosslinking studies to determine the potential ligand binding sites. While the molecular structure of GPRC5A has not been determined, RosettaGPCR^49^, AlphaFold^50^ and Robetta^51^ provide structural models of its 7-helix transmembrane domain (**Fig. 6a, Supplementary Fig. 8a**). Based on the knowledge of ligand binding to aminergic GPCRs and class C GPCRs, we mutated a predicted extracellular patch of acidic amino acid residues as well as a polar and aromatic pocket within the transmembrane domain (**Fig. 6a**). Using the PRESTO-Tango assay upon 7FTA treatment, we found that the extracellular patch of aspartic acids in GPRC5A are not involved in the interaction with aromatic monoamines (**Fig. 6b**). Mutations of N252 and F256 respectively to alanine resulted in decreased GPRC5A activation with 7FTA, which indicates that these two amino acids within a polar and aromatic pocket of transmembrane domain may be crucial for ligand binding (**Fig. 6b**). Alanine mutations of F105, F109, S220 and W224 had no significant effect on 7FTA activation of GPRC5A (**Fig. 6b**). We then generated the N252A and F256A double mutants and found that both single and double mutations decreased the 7FTA mediated GPRC5A activation (**Fig. 6c**) and photoaffinity labeling (**Fig. 6d,e**). These results suggest N252 and F256 are key amino acid residues involved in GPRC5A ligand binding. Nonetheless, these amino acid residues are conserved amongst retinoic acid-induced class C orphan GPCRs (GPRC5A, GPRC5B, GPRC5C and GPRC5D) (**Supplementary Fig. 8b**), which are all activated by 7FTA (**Fig. 6f**). Moreover, both TA and 7FTA function as agonists of GPCRs sensing aromatic monoamines, such as ADRA2A, DRD2 and HTR4 (**Fig. 6g**). However, GPRC5A did not respond to endogenous neurotransmitters like epinephrine, dopamine and serotonin (**Fig. 6h**), indicating distinct mechanisms and functions of metabolite sensing compared to several canonical GPCRs. Collectively, our results demonstrate that aromatic amines can bind a conserved transmembrane domain of GPRC5A and activate other retinoic acid-induced class C GPCRs.

**Fig. 6.**
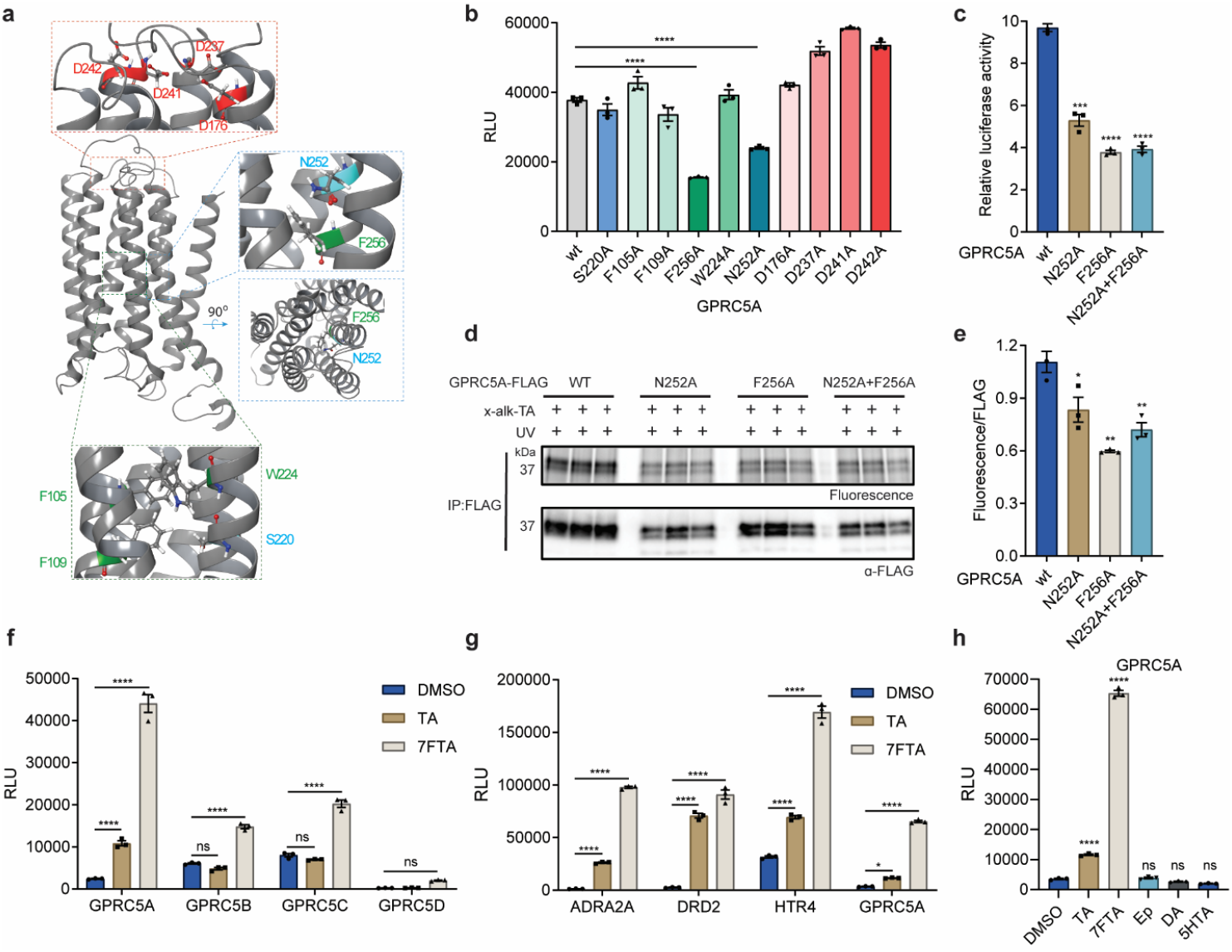
Analysis of aromatic monoamine-GPRC5A interactions. **a**, Molecular model of GPRC5A made from RosettaGPCR. **b-c**, PRESTO-Tango assay for mutagenesis screening of key residues in GPRC5A-7FTA (100 μM) interactions. **d-e**, Photoaffinity crosslinking of x-alk-TA to FLAG-tagged GPRC5A mutants in HEK293T cells revealed by rhodamine labeling. **f-g**, Evaluation of 100 μM of TA and 7FTA on class C orphan GPCRs (GPRC5A,B,C,D) and aminergic GPCRs (ADRA2A, DRD2, HTR4) activation. **h**, Examination of endogenous neurotransmitters epinephrine (10 μM), dopamine (10 μM) and serotonin (100 μM) on GPRC5A activation. Data indicates mean with SEM from three replicates. One-way ANOVA using Dunnett’s multiple comparisons test (b,c,e,h) or Two-way ANOVA using Tukey’s multiple comparisons test (f,g). *P<0.05, **P<□0.01, ***P≤□0.001 ****P≤□0.0001, ns indicates not significant.

## DISCUSSION

The human microbiota and their metabolites are important modulators of host physiology. Understanding the molecular basis by which specific microbiota species and metabolites exert their mechanisms of action is therefore crucial for dissecting hostmicrobe interactions. For example, aAAs are metabolized by specific microbiota into diverse molecules that play pivotal roles in host health and disease.^14,52^ Desaminotyrosine (DAT) produced by *Clostridium orbiscindens* protects mice from influenza virus challenge through augmentation of type I IFN signaling and suppression of lung immunopathology.^53^ Alternatively, *Lactobacillus reuteri* generates indole-3-lactic acid (ILA) and IAld that activates AhR and induces intraepithelial CD4+ CD8αα+ T cells to maintain intestinal homeostasis.^54^ Moreover, indole-3-ethanol, indole-3-pyruvate and IAld can regulate gut barrier function through AhR.^55^ Indole metabolites such as IAA and IPA have also been reported to activate pregnane X receptor (PXR) and increases tight junction protein expression and downregulate the expression of pro-inflammatory cytokines.^17^ Beyond modulating intestinal homeostasis, *C. sporogenes* has been reported to metabolize tryptophan and generate high levels of indole metabolites that can be detected in serum and modulate systemic immune responses.^15,56^ While certain protein targets and receptors have been reported for these indole metabolites, the relatively high levels (micro-millimolar levels) of these metabolites in the intestinal lumen and circulation, suggests they may engage multiple protein targets in host cells.

To address this general challenge of investigating microbiota metabolite mechanisms, we employed chemical proteomics and explored the protein targets of IAA, a prominent microbiota metabolite^14,15^ with potentially multiple mechanisms of action^16,17^. Through the generation of metabolic and photoaffinity IAA-based chemical reporters, we identified many indole-metabolite interacting proteins. We focused on GPRC5A and discovered that this previously annotated class C orphan GPCR is able to bind indole metabolites, but is only activated by aromatic monoamines such as TA, PEA or tyramine. These exciting discoveries reveal the first small molecule agonists of GPRC5A and its related class C GPCR homologs (GPRC5B, GPRC5C and GPRC5D). Moreover, this led us to identify and characterize aromatic monoamine-producing microbiota species (*R. gnavus, M. morganii* and *Enterococcus* species) and their associated decarboxylases that can generate aromatic amines and activate GPRC5A signaling (**Supplementary Fig. 9**). In addition, our analysis of synthetic aromatic amines revealed 7FTA as a more potent agonist of GPRC5A and its homologs as well as potential sites of ligand binding within their 7-transmembrane domains. Of note, our proteomic studies of IAA-interacting proteins also identified additional candidate GPCRs (GPR107 and GPR108) that have no reported small molecule ligands and numerous other proteins, which may reveal additional mechanisms of action for IAA. Chemoproteomics therefore provides a complementary approach to targeted screening of individual microbiota species^19,57,58^ or biosynthetic gene clusters^19,57,58^ for novel GPCR ligands.

Our discovery of microbiota metabolites and synthetic ligands of GPRC5A is important, as GPRC5A expression is correlated with lung, pancreatic and colon cancers^30,31,59–61^. Notably, *Gprc5a−/−* mice exhibit higher inflammation signatures and develop lung cancer over time^29,32^. In both human and mouse studies, GPRC5A has been implicated in epidermal growth factor receptor (EGFR)^62^ and NF-κB^32^ signaling, which may account for the modulation of inflammation and cancer incidence. However, no small-molecule agonists and antagonists have been reported for GPRC5A or its homologs. Our results suggest the generation of aromatic monoamines by specific microbiota species are sensed by GPRC5A, which may crosstalk with cell signaling of other monoamine-sensing GPCRs (ADRA2A, DRD2 and HTR4) and/or key pathways associated with tissue homeostasis or immunity (**Supplementary Fig. 9**). Therefore, the future development of more potent and specific GPRC5A agonists and antagonists based on our studies may provide important pharmacological tools and therapeutic leads to investigate the roles of GPRC5A in physiology and human disease. Lastly, our studies highlight the utility of chemoproteomics to undercover novel mechanisms of action for indole metabolites that could be applied to other classes of microbiota metabolites.

## Acknowledgements

We thank H. Molina from Proteomics Resource Center at The Rockefeller University for proteomic analysis. We also thank G. Barnea (Brown University) for providing HTLA cells, B. Roth (UNC) for PRESTO-Tango plasmids, M. Gilmore (Harvard Medical School) for *E. faecium* Com15, S. F. Brady (The Rockefeller University) for *R. gnavus* ATCC 29149, and C-J. Guo (Weill Cornell Medical School) for *C. sporogenes* ATCC 15579. K.S. acknowledges support from Scripps Research Department of Immunology and Microbiology T32AI007244. H.C.H. acknowledges grant support from NIH-NIGMS R01GM087544.

## Author contributions

X.Z. and H.C.H. conceived of the project. K.R.S. contributed to the analysis of GPRC5A signaling, V.C. prepared TyrDC-KO *E. faecium* Com15 and M.E.G. performed phylogenetic analysis of bacterial decarboxylases. X.Z. performed the experiments and interpreted the data. X.Z. and H.C.H. wrote the manuscript with input of editing from K.R.S., V.C. and M.E.G.

## Additional information

Supplementary information is available for this paper.

---

**Supplementary Fig. 1.**
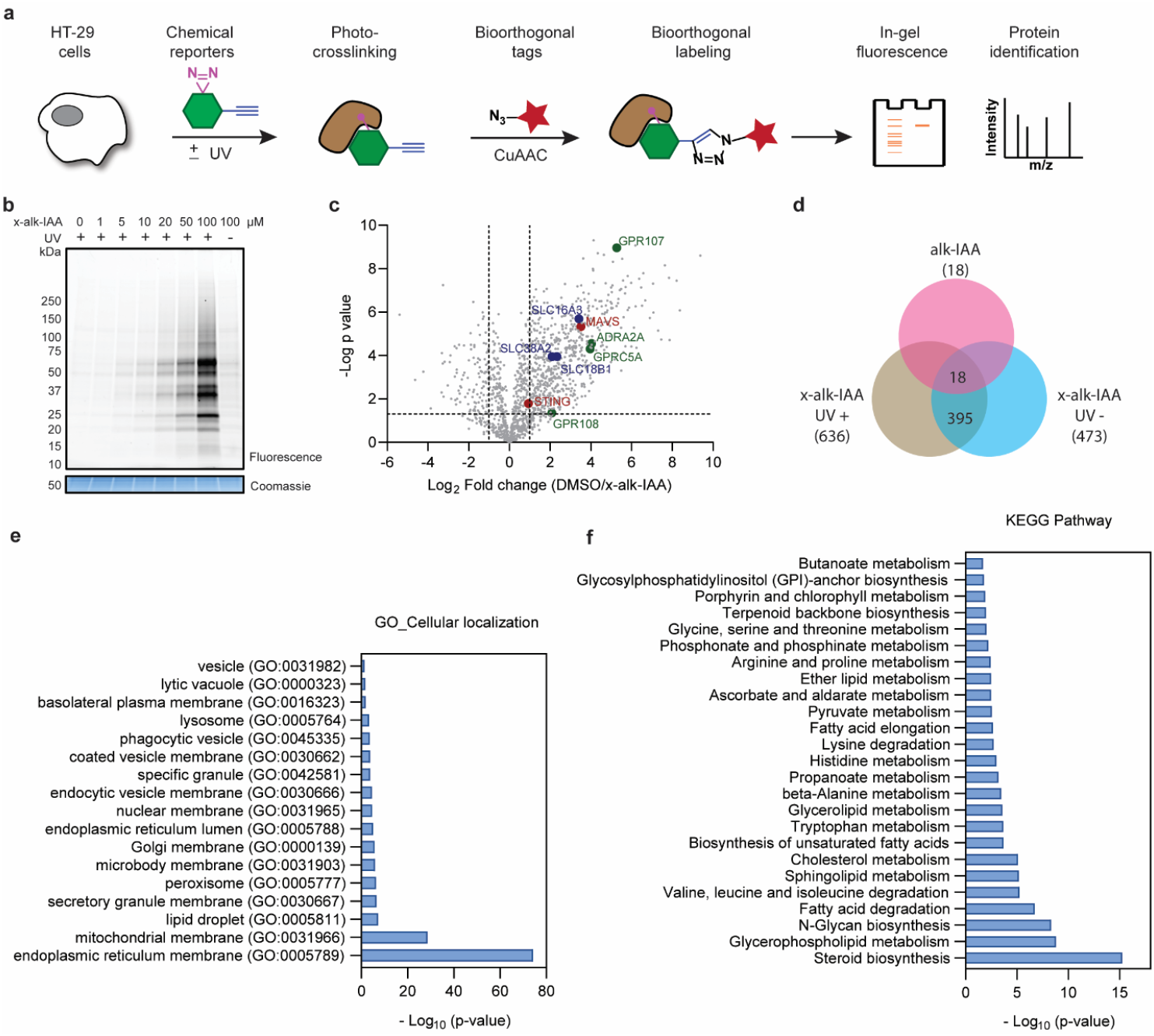
Chemoproteomic analysis of IAA-interacting proteins in HT-29 cells. **a**, Scheme for in-gel fluorescence profiling and chemoproteomic analysis of microbiota metabolite protein targets in HT-29 cells using photoaffinity chemical reporters. **b**, In-gel fluorescence profiling of IAA-interacting proteins in HT-29 cells using indicated concentration of x-alk-IAA with or without UV irradiation. **c**, Volcano plot of the identified proteins from HT-29 cells by chemical proteomics using x-alk-IAA with UV irradiation compared to DMSO control (p value is 0.05, two folds change in four replicates). Some of key hits are highlighted in GPCRs (green), transporters (blue) and immunity-associated proteins (red). **d**, Venn diagram for the statistically significant hits identified by chemical proteomics using two chemical reporters with or without UV irradiation. **e**, Gene ontology analysis of the major subcellular localization of x-alk-IAA interacting proteins. **f**, Major KEGG pathways involve x-alk-IAA interacting proteins.

**Supplementary Fig. 2.**
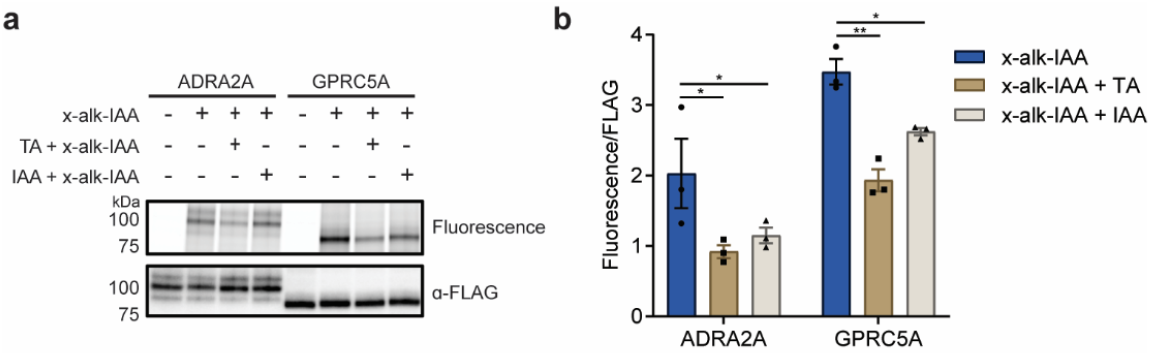
ADRA2A and GPRC5A are crosslinked by IAA photoaffinity chemical reporters. **a,b,** Competitive labeling assay using 200 μM IAA or TA that compete with 20 μM x-alk-IAA in binding on ADRA2A or GPRC5A. Two-way ANOVA using Holm-Sidak’s multiple comparisons test. *P<0.05, **P≤□0.01, ns indicates not significant.

**Supplementary Fig. 3.**
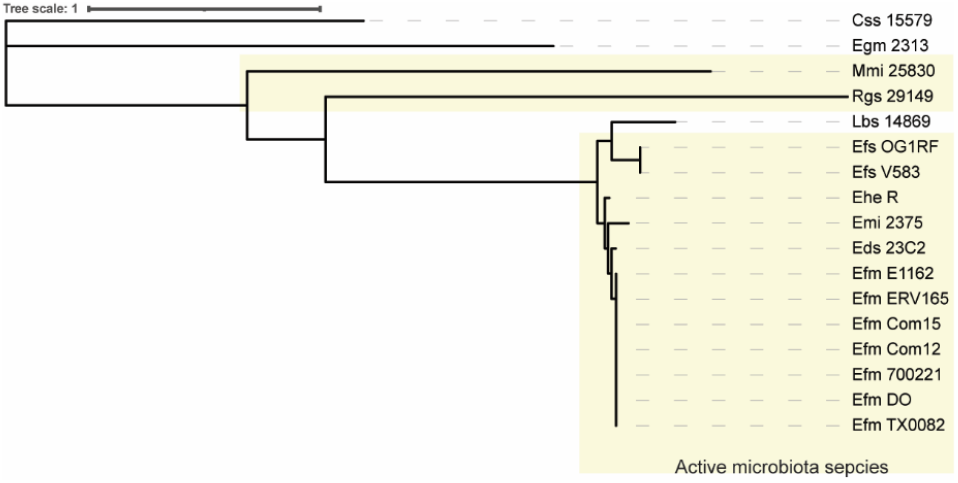
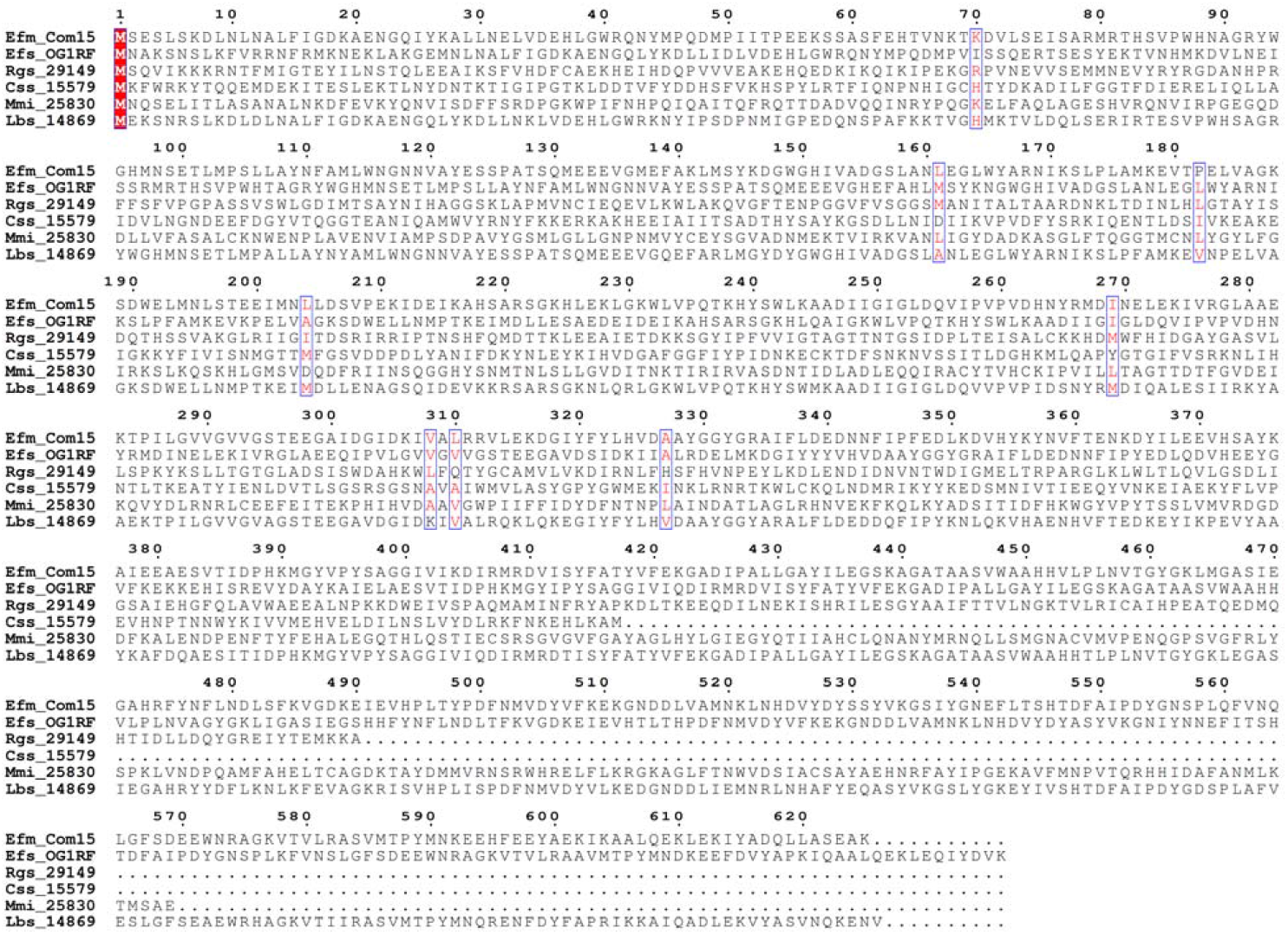
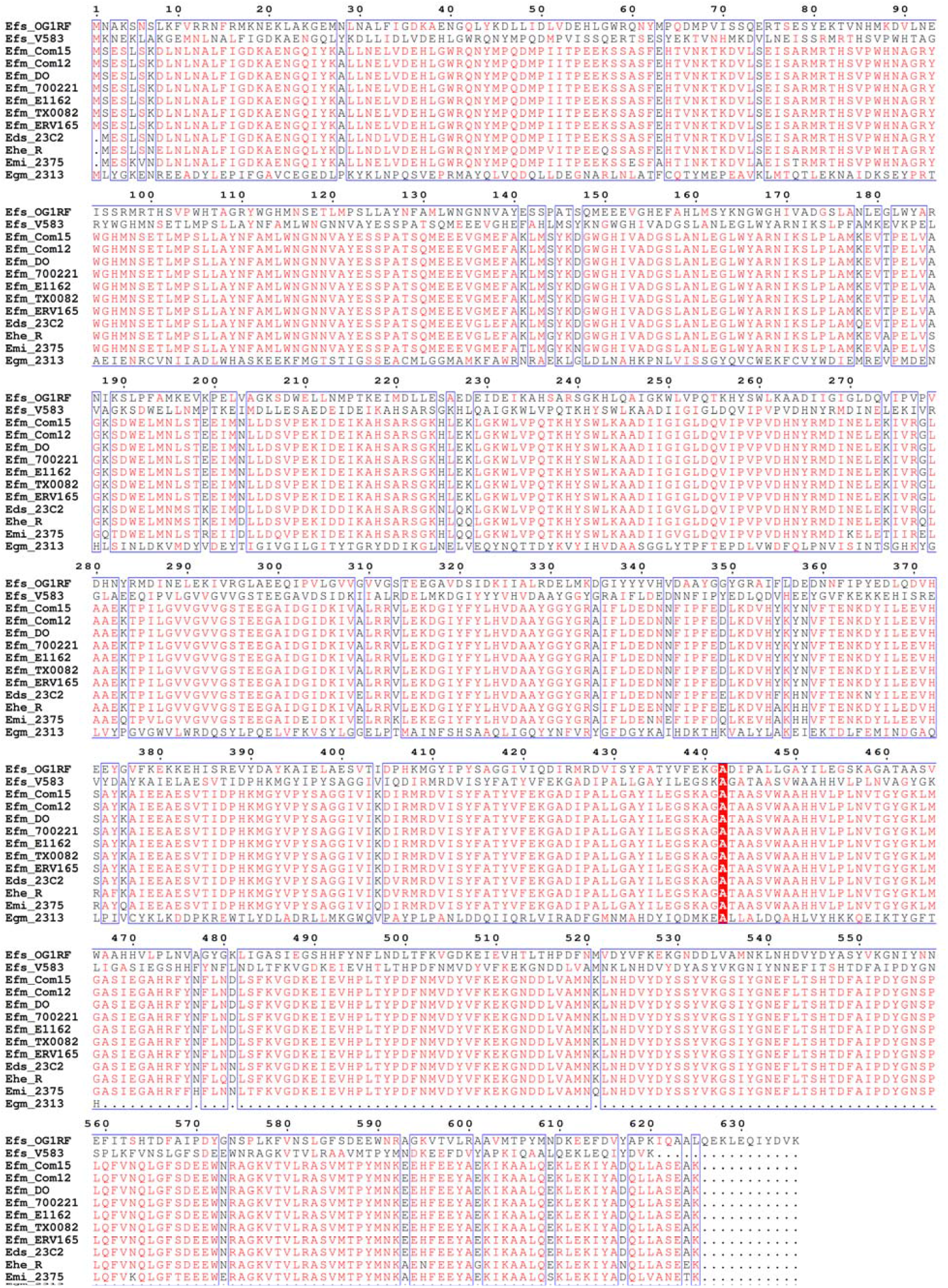
**a**, Unrooted phylogenetic clustering of protein sequences of tyrosine decarboxylases of *Enterococcus* species and strains, as well as *L. brevis* 14869, glutamate decarboxylase of *E. gallinarum* 2313, tryptophan decarboxylases of *R. gnavus* 29149 and *C. sporogenes* 15579, glutamate or tyrosine decarboxylases of *M. morganii* 25830. Active strains are indicated by the yellow box. Scale bar represents sequence distance. **b,** Sequence alignment of amino acid decarboxylases of different microbiota species. **c**, Sequence alignment of amino acid decarboxylases of different *Enterococcus* species.

**Supplementary Fig. 4.**
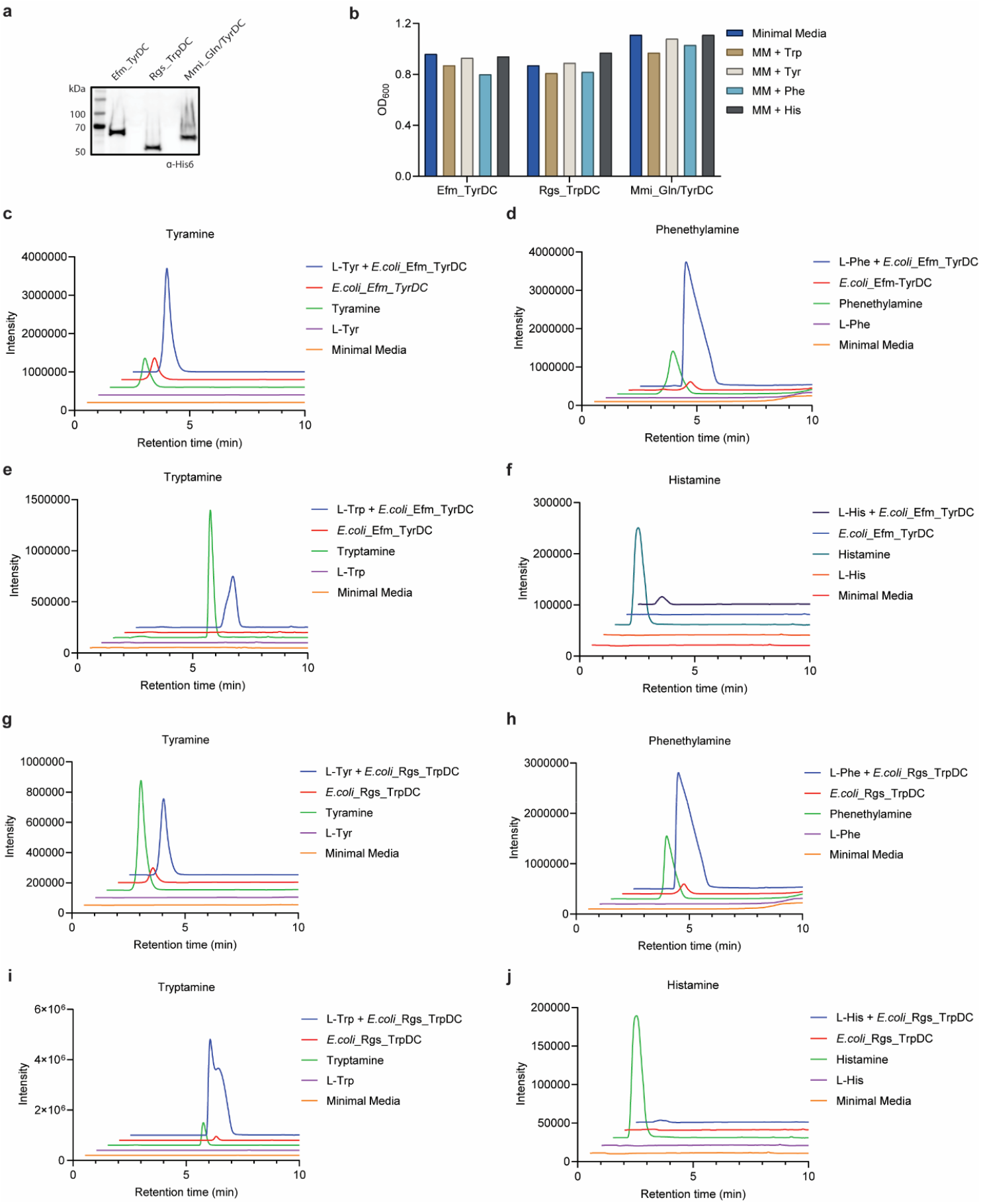
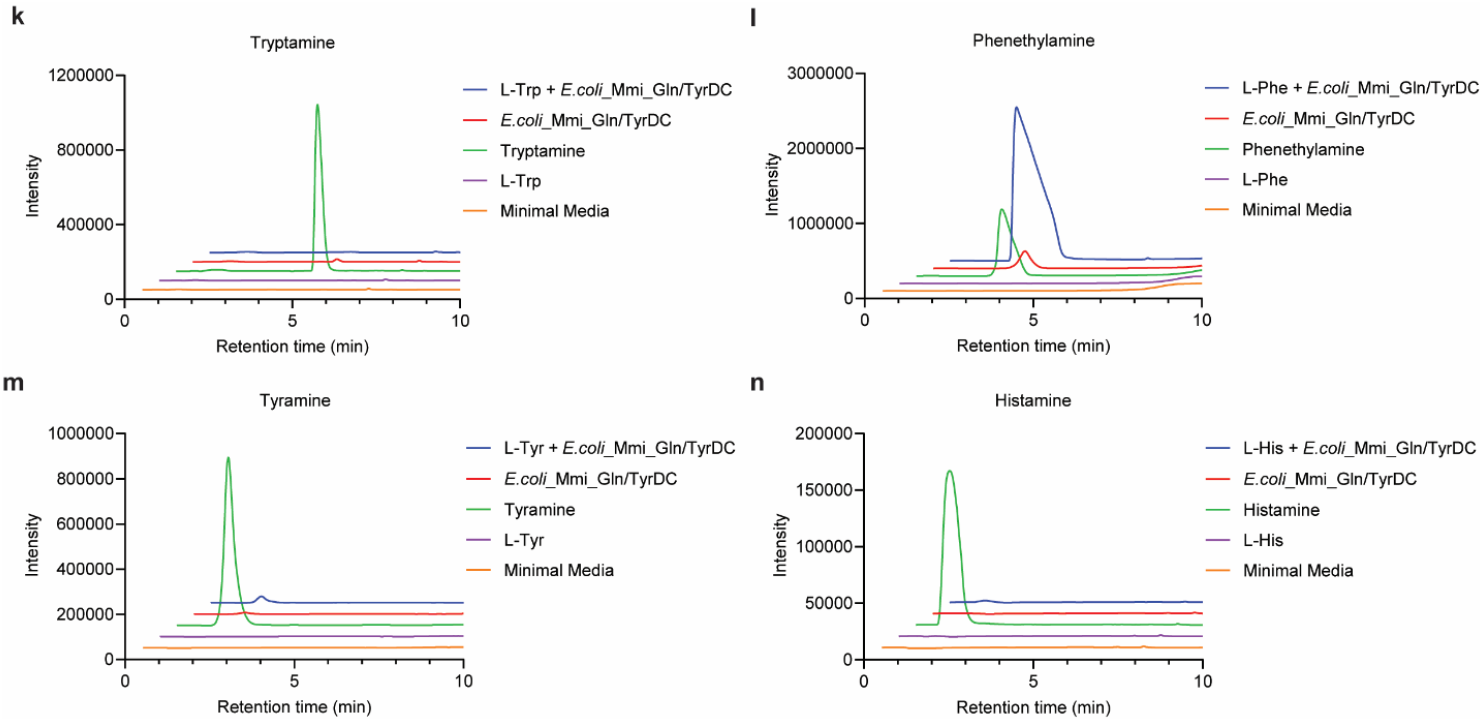
Bacterial decarboxylases transform specific aAAs to aromatic monoamines. **a**, Western blot analysis of the overexpression of His6-tagged *E. faecium* Com15 tyrosine decarboxylase (Efm_TyrDC), *R. gnavus* tryptophan decarboxylase (Rgs_TrpDC) and *M. morganii* glutamate/tyrosine decarboxylase (Mmi_Gln/TyrDC) in *E. coli* DH5a. **b**, Bacterial growth in minimal media supplemented with individual aAAs at 37 °C for 16 h. **c-n**, LC-MS analysis of respective aromatic monoamines produced by *E. coli*-Efm_TyrDC (c-f), *E. coli*_Rgs_TrpDC (g-j) and *E. coli*_ Mmi_Gln/TyrDC (k-n).

**Supplementary Fig. 5.**
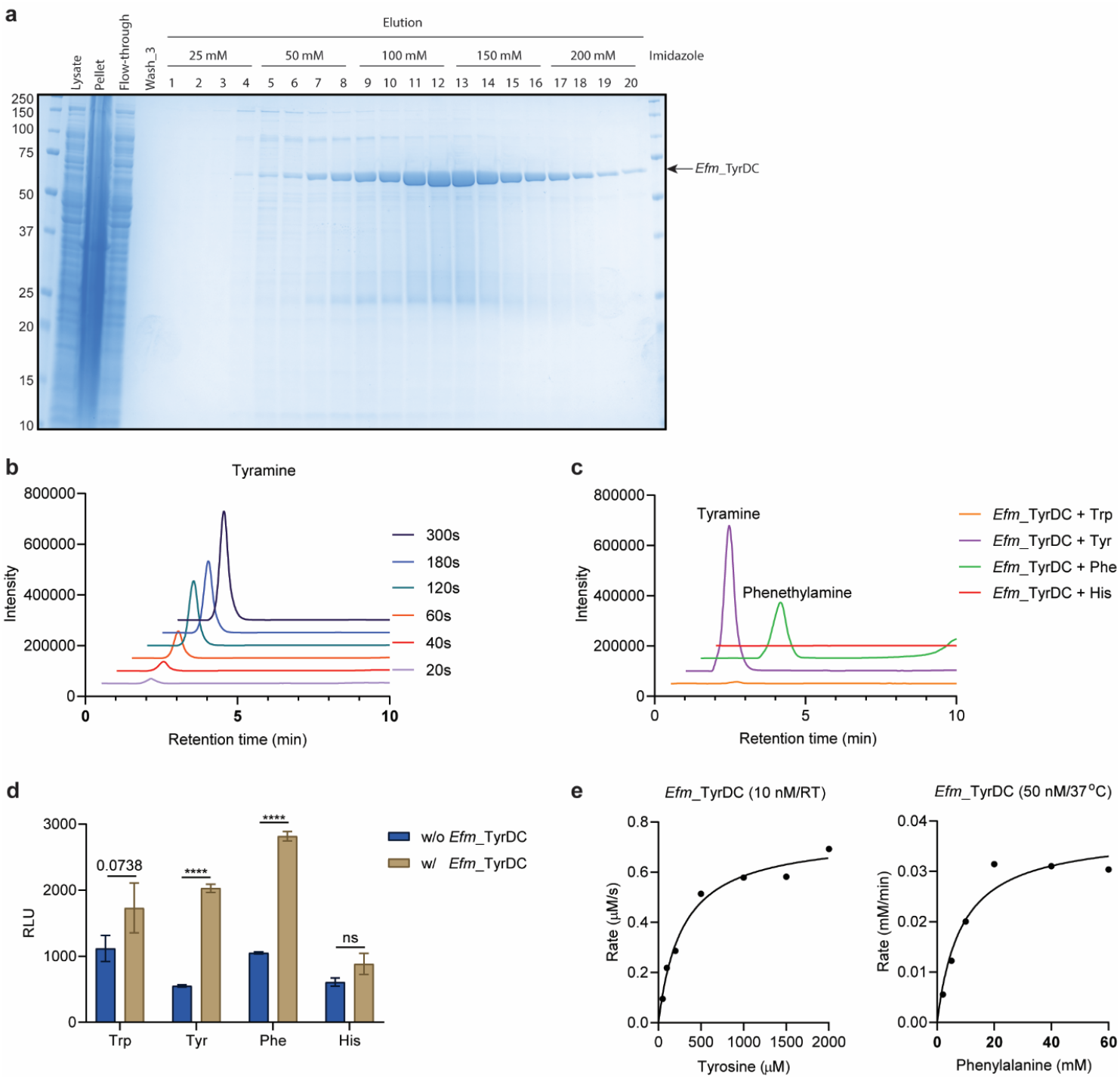
*In vitro* enzymatic transformation of aAAs to aromatic monoamines. **a**, SDS-PAGE analysis of the purification of His6-tagged *E. faecium* Com15 tyrosine decarboxylase (Efm_TyrDC). **b**, LC-MS analysis of the time-dependent formation of tyramine from enzymatic transformation of tyrosine by 100 nM Efm_TyrDC at room temperature. **c**, LC-MS determination of tryptamine, tyramine, phenethylamine and histamine from enzymatic transformation of tryptamine, tyrosine, phenylalanine and histidine by 100 nM Efm_TyrDC at 37 °C for 10 min. **d**, PRESTO-Tango assay for Efm_TyrDC enzymatic products for GPRC5A activation. **e**, Michaeslis-Menten kinetics study of *Efm*_TyrDC catalyzed transformation of tyrosine and phenylalanine. Data indicates mean with SEM from three replicates. Twoway ANOVA using Sidak’s multiple comparisons test. ****p<0.0001, ns indicates not significant.

**Supplementary Fig. 6.**
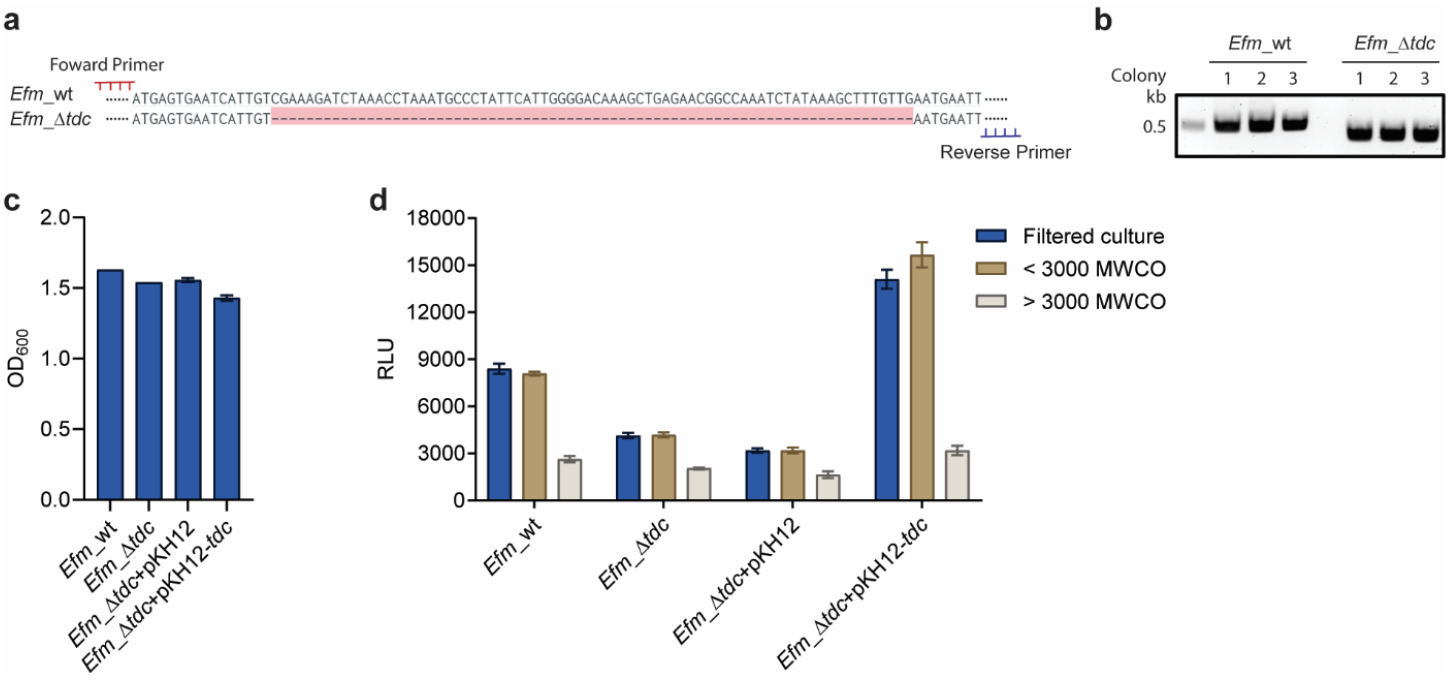
Knock-out of tyrosine decarboxylase in *E. faecium* Com15. **a**, Genetic deletion of segment of DNA disrupts *TyrDC* gene expression in *E. faecium* Com15. **b**, Agarose gel electrophoresis of the PCR products from wild-type and *TyrDC* gene-disrupted *E. faecium* Com15. **c**, Optical density measurement of bacterial growth including wild-type *E. faecium* Com15 (Efm_wt), *E. faecium* Com15 *TyrDC* knock-out (Efm_Δtdc), complementation of vector control (pKH12) or TyrDC (pKH12-tdc) in Efm_Δtdc at 37 °C for 16 h. **d**, PRESTO-Tango assay for examining bacterial cultures for GPRC5A activation. Bacterial growth media were filtered through 0.2 uM membrane followed by 3,000 MWKO membrane.

**Supplementary Fig. 7.**
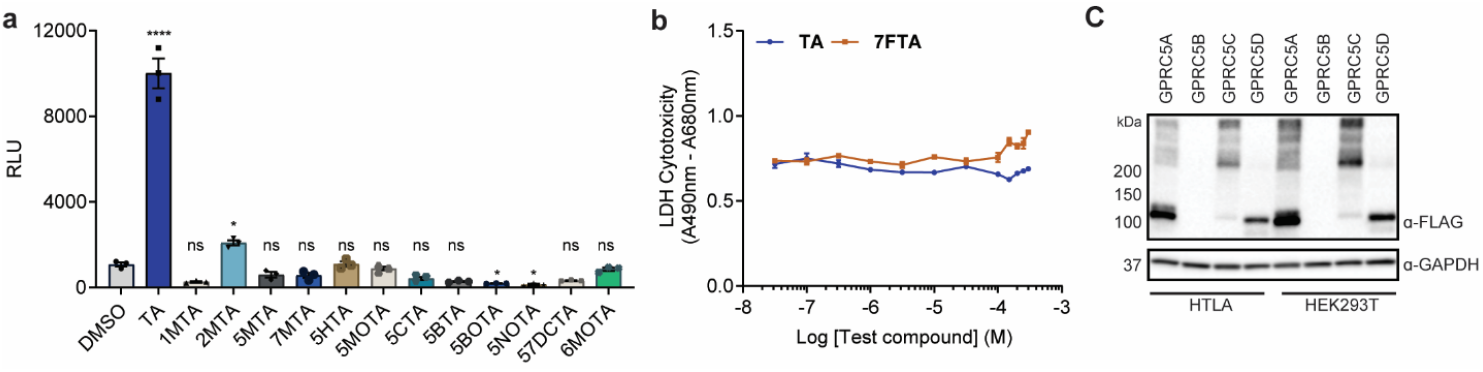
Structure-activity relationship studies of GPRC5A ligands. **a**, PRESTO-Tango assay for screening a collection of tryptamine derivatives (100 μM) for GPRC5A activation. **b**, LDH cytotoxicity assay to examine TA and 7FTA in HEK293T cells. **c**, Western blot analysis of the expression of GPRC5A-, GPRC5B-, GPRC5C-, GPRC5D-Tango proteins in HTLA or HEK293T cells, respectively.

**Supplementary Fig. 8.**
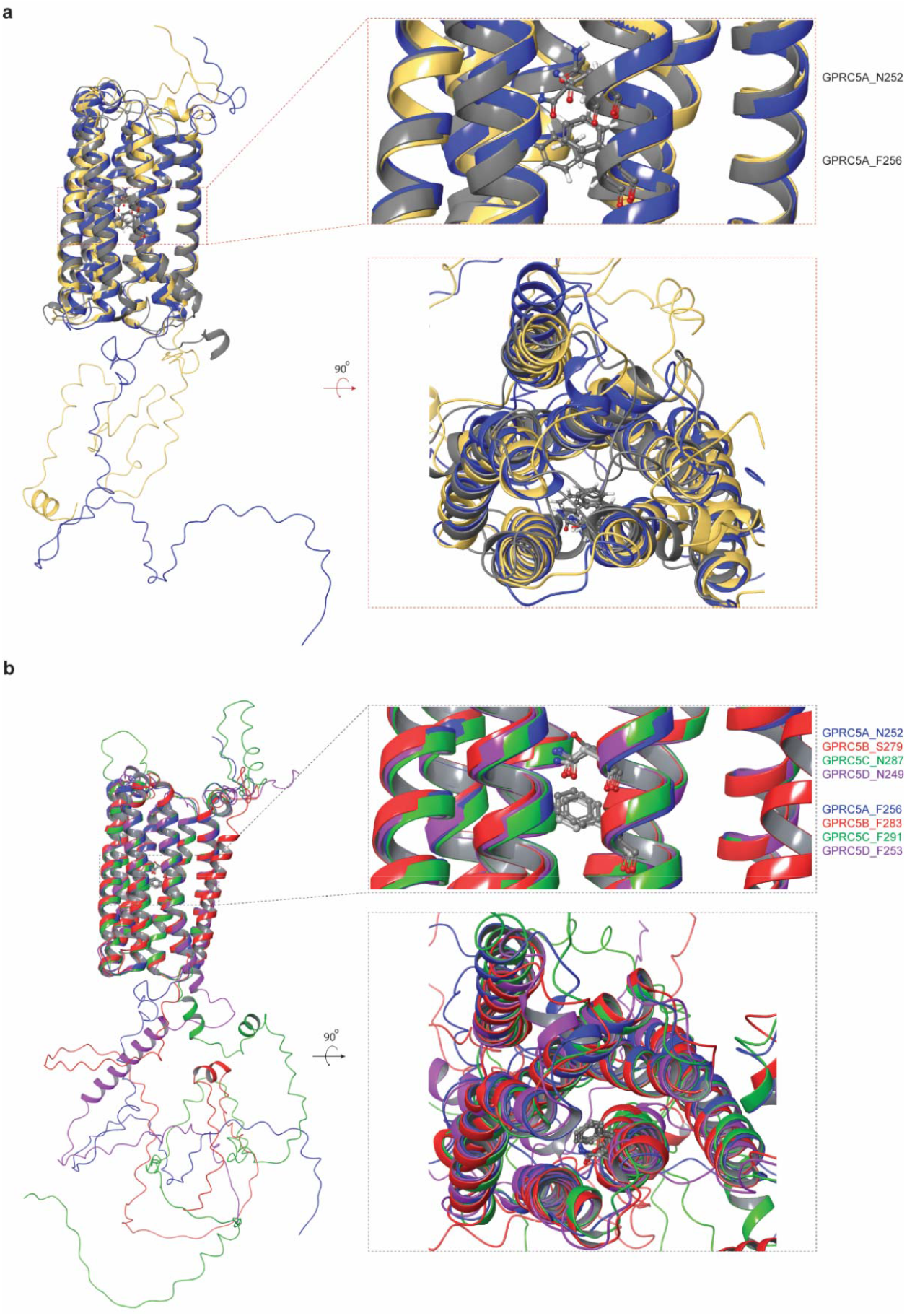
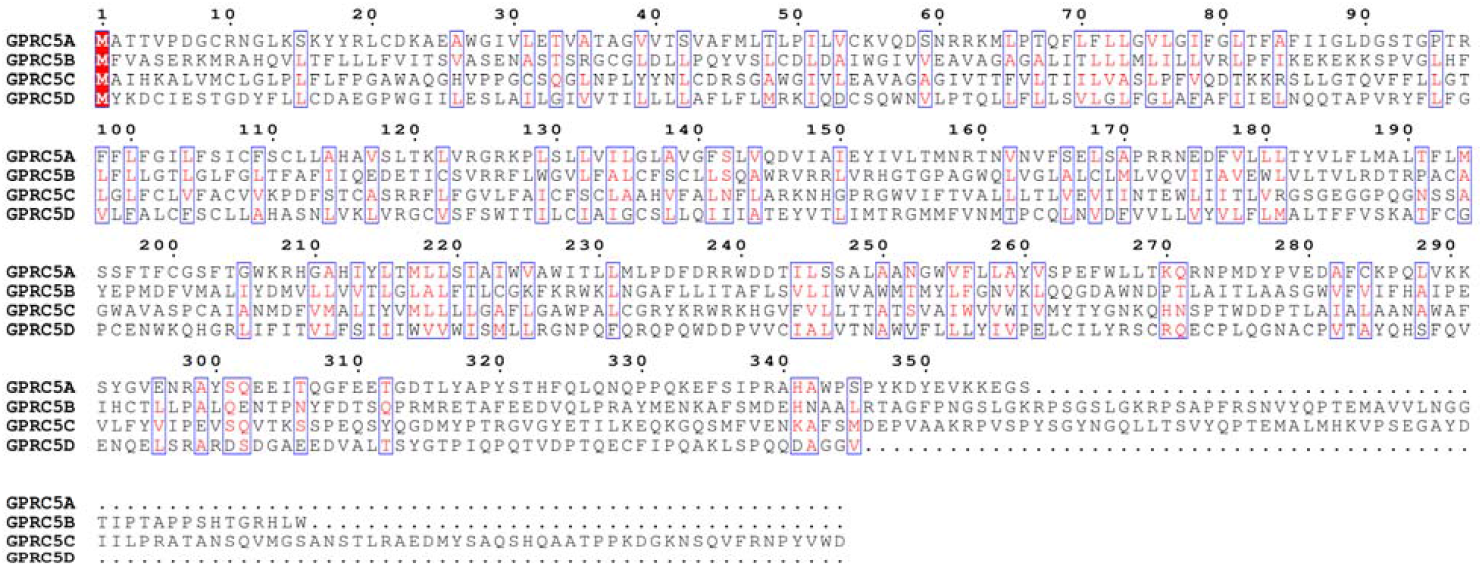
**a,** Structural models of GPRC5A from RosettaGPCR (gray), AlphaFold(blue) and Robetta (yellow). N252 and F256 are indicated. **b**, Structural models of GPRC5A (blue), GPRC5B (red), GPRC5C (green) and GPRC5D (purple) from AlphaFold. **c,** Sequence alignment of GPRC5A, GPRC5B, GPRC5C, GPRC5D.

**Supplementary Fig. 9.**
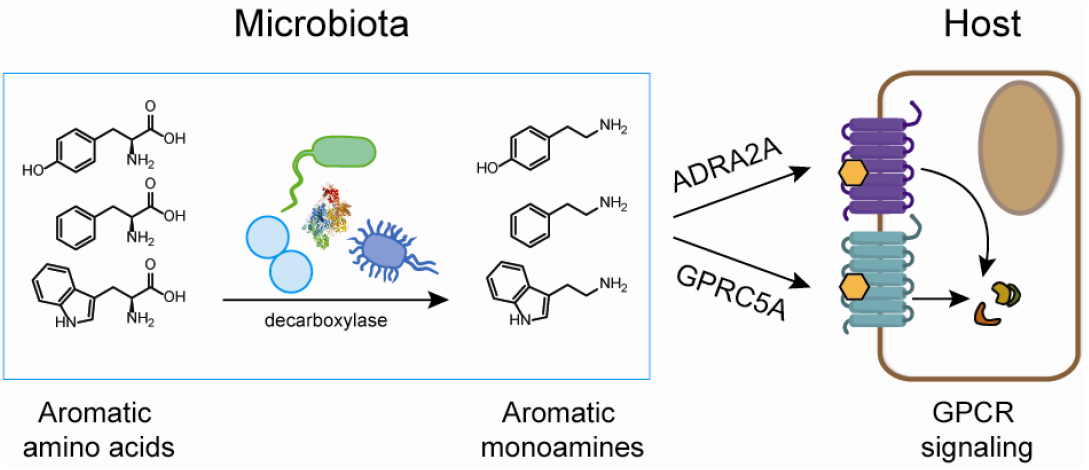
Schematic of specific microbiota metabolism of aromatic amino acids by decarboxylases and activation of GPRC5A and other functionally-related GPCRs within host cells.

**Supplementary Table 1.**
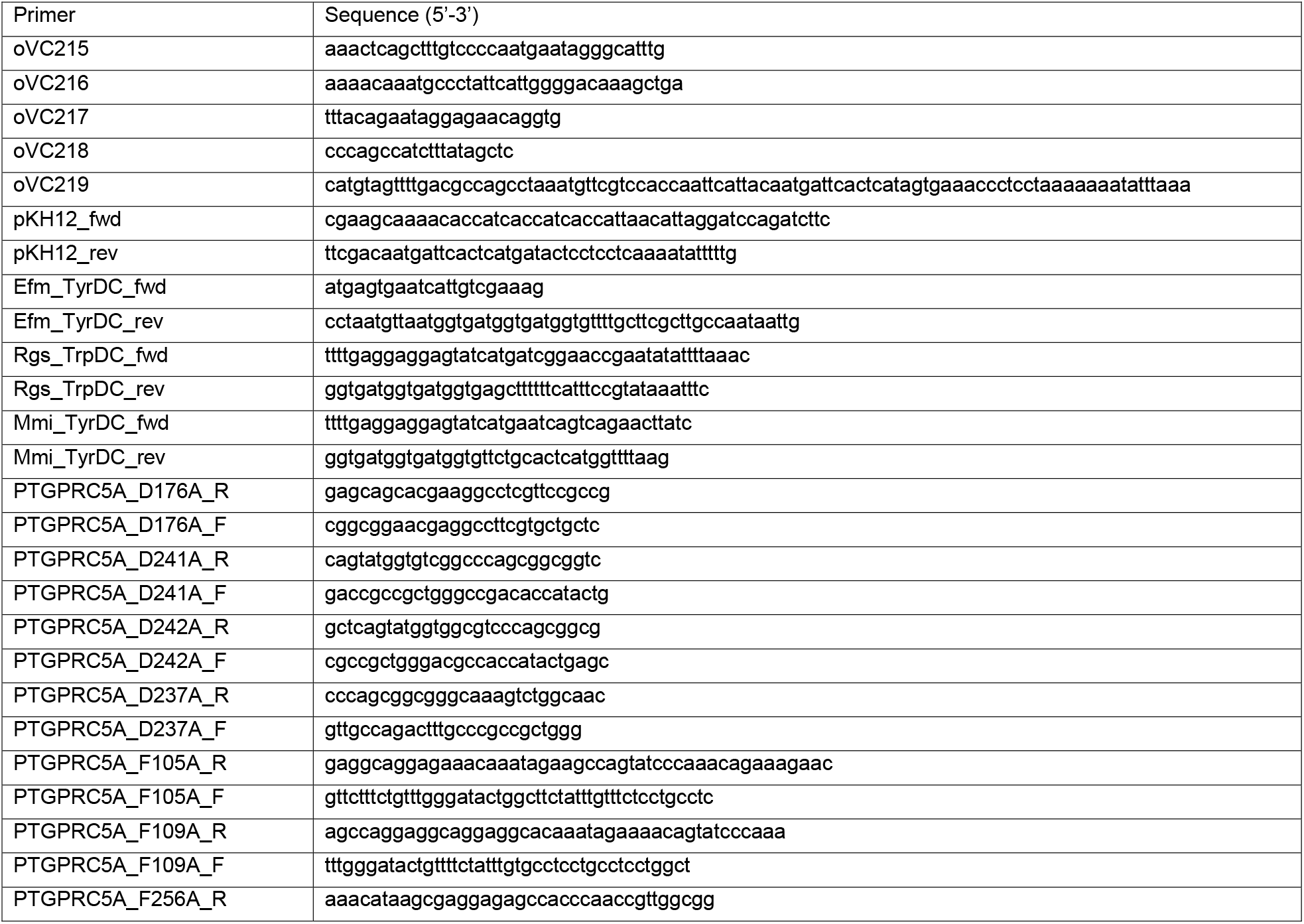
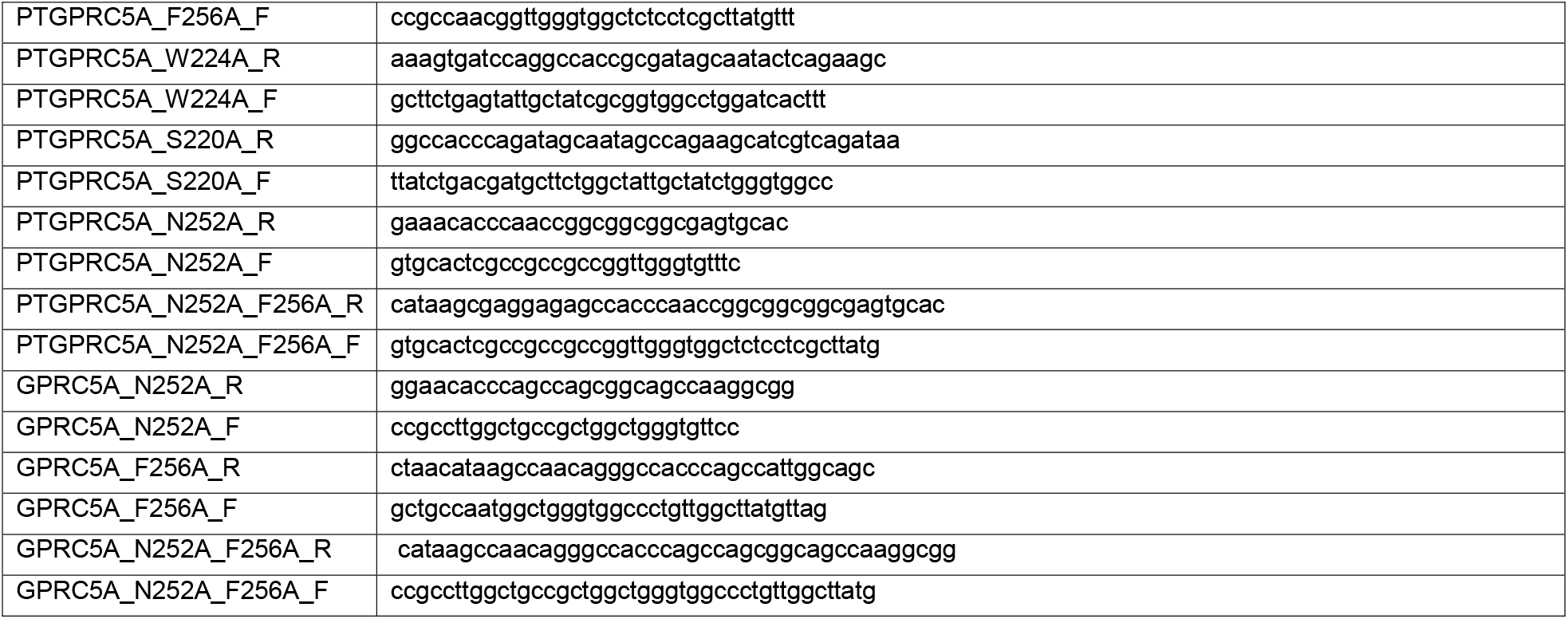
Primers.

**Supplementary Table 2.** Plasmids.

| Plamid | Gene | Antibiotic resistance | Origin |
| --- | --- | --- | --- |
| pRecT | recT | Erythromycin | Our lab |
| pCas9-Efm_TyrDC | Cas9/sgRNA | Chloramphenicol | This study |
| pKH12-Efm_TyrDC | Efm_TyrDC-His6 | Chloramphenicol | This study |
| pKH12-Rgs_TrpDC | Rgs_TrpDC-His6 | Chloramphenicol | This study |
| pKH12-Mmi_Gln/TyrDC | Mmi_Gln/TyrDC-His6 | Chloramphenicol | This study |
| pGL4.43_Luc2P/XRE/Hygro | 3x XRE-Luc | Ampicillin | Promega E4121 |
| pRL-TK | Renilla-Luc | Ampicillin | Promega E2241 |
| pCMV3-N-HA-AhR | HA-AhR | Kanamycin | SinoBio HG10456-NY |
| pcDNA3.1_ADRA2A-Tango | Flag-ADRA2A-tTA | Ampicillin | Addgene 66216 |
| pcDNA3.1_DRD2-Tango | Flag-DRD2-tTA | Ampicillin | Addgene 66269 |
| pcDNA3.1_HTR4-Tango | Flag-HTR4-tTA | Ampicillin | Addgene 66412 |
| pcDNA3.1_GPRC5D-Tango | Flag-GPRC5D-tTA | Ampicillin | Addgene 66385 |
| pcDNA3.1_GPRC5C-Tango | Flag-GPRC5C-tTA | Ampicillin | Addgene 66384 |
| pcDNA3.1_GPRC5B-Tango | Flag-GPRC5B-tTA | Ampicillin | Addgene 66383 |
| pcDNA3.1_GPRC5A-Tango | Flag-GPRC5A-tTA | Ampicillin | Addgene 66382 |
| pcDNA3.1_GPRC5A_D176A-Tango | Flag-GPRC5A_D176A-tTA | Ampicillin | This study |
| pcDNA3.1_GPRC5A_D241A-Tango | Flag-GPRC5A_D241A-tTA | Ampicillin | This study |
| pcDNA3.1_GPRC5A_D242A-Tango | Flag-GPRC5A_D242A-tTA | Ampicillin | This study |
| pcDNA3.1_GPRC5A_D237A-Tango | Flag-GPRC5A_D237A-tTA | Ampicillin | This study |
| pcDNA3.1_GPRC5A_F105A-Tango | Flag-GPRC5A_F105A-tTA | Ampicillin | This study |
| pcDNA3.1_GPRC5A_F109A-Tango | Flag-GPRC5A_F109A-tTA | Ampicillin | This study |
| pcDNA3.1_GPRC5A_F256A-Tango | Flag-GPRC5A_F256A-tTA | Ampicillin | This study |
| pcDNA3.1_GPRC5A_W224A-Tango | Flag-GPRC5A_W224A-tTA | Ampicillin | This study |
| pcDNA3.1_GPRC5A_S220A-Tango | Flag-GPRC5A_S220A-tTA | Ampicillin | This study |
| pcDNA3.1_GPRC5A_N252A-Tango | Flag-GPRC5A_N252A-tTA | Ampicillin | This study |
| pcDNA3.1_GPRC5A_N252A_F256A-Tango | Flag-GPRC5A_N252A_F256A-tTA | Ampicillin | This study |
| pCMV3-GPRC5A | GPRC5A | Ampicillin | SinoBio HG12833-UT |
| pcDNA3_c-Flag | Flag | Ampicillin | Addgene 20011 |
| pcDNA3_GPRC5A-Flag | GPRC5A-Flag | Ampicillin | This study |
| pcDNA3_GPRC5A_N252A-Flag | GPRC5A_N252A-Flag | Ampicillin | This study |
| pcDNA3_GPRC5A_F256A-Flag | GPRC5A_F256A-Flag | Ampicillin | This study |
| pcDNA3_GPRC5A_N252A_F256A-Flag | GPRC5A_N252A_F256A-Flag | Ampicillin | This study |

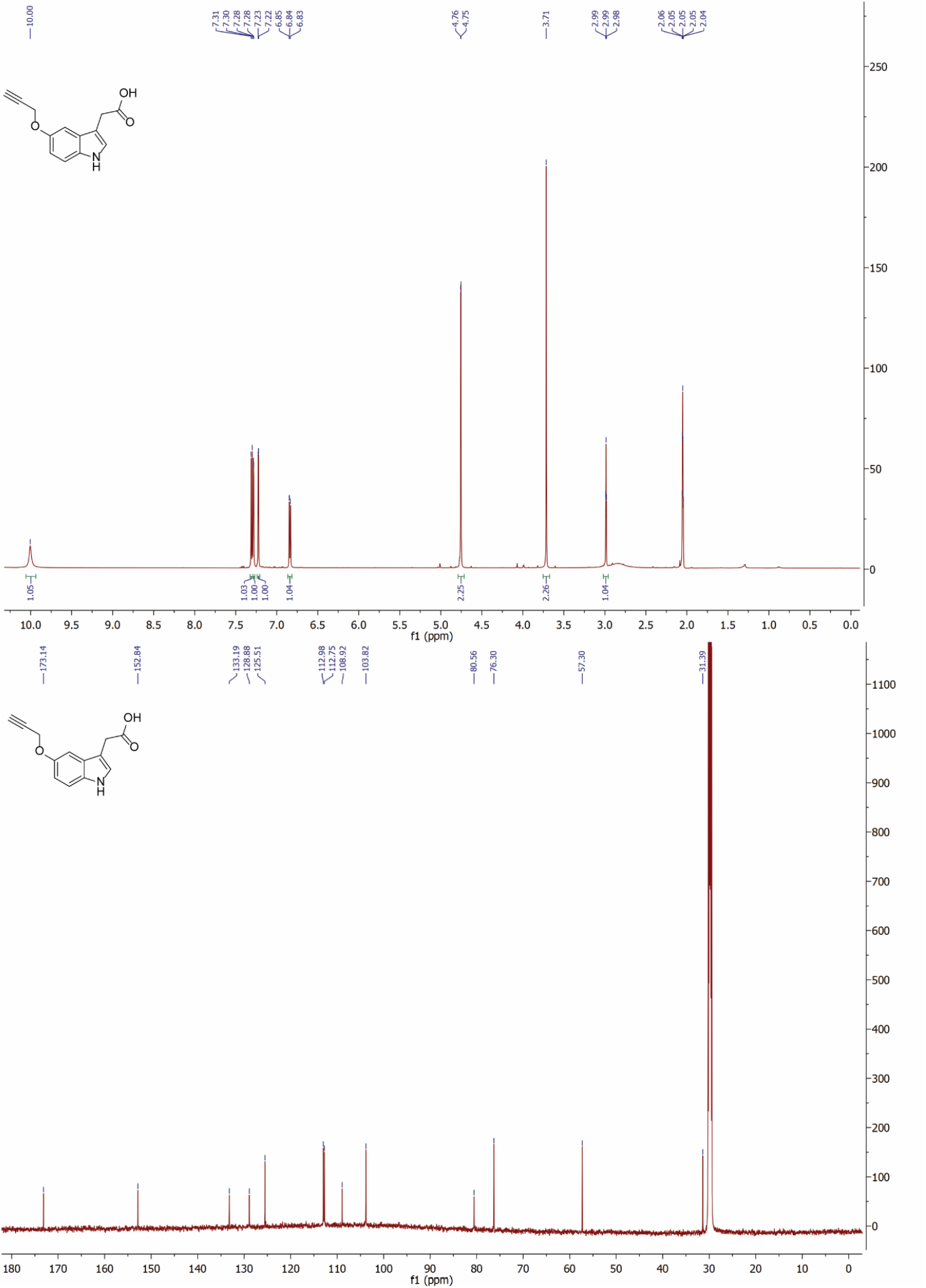

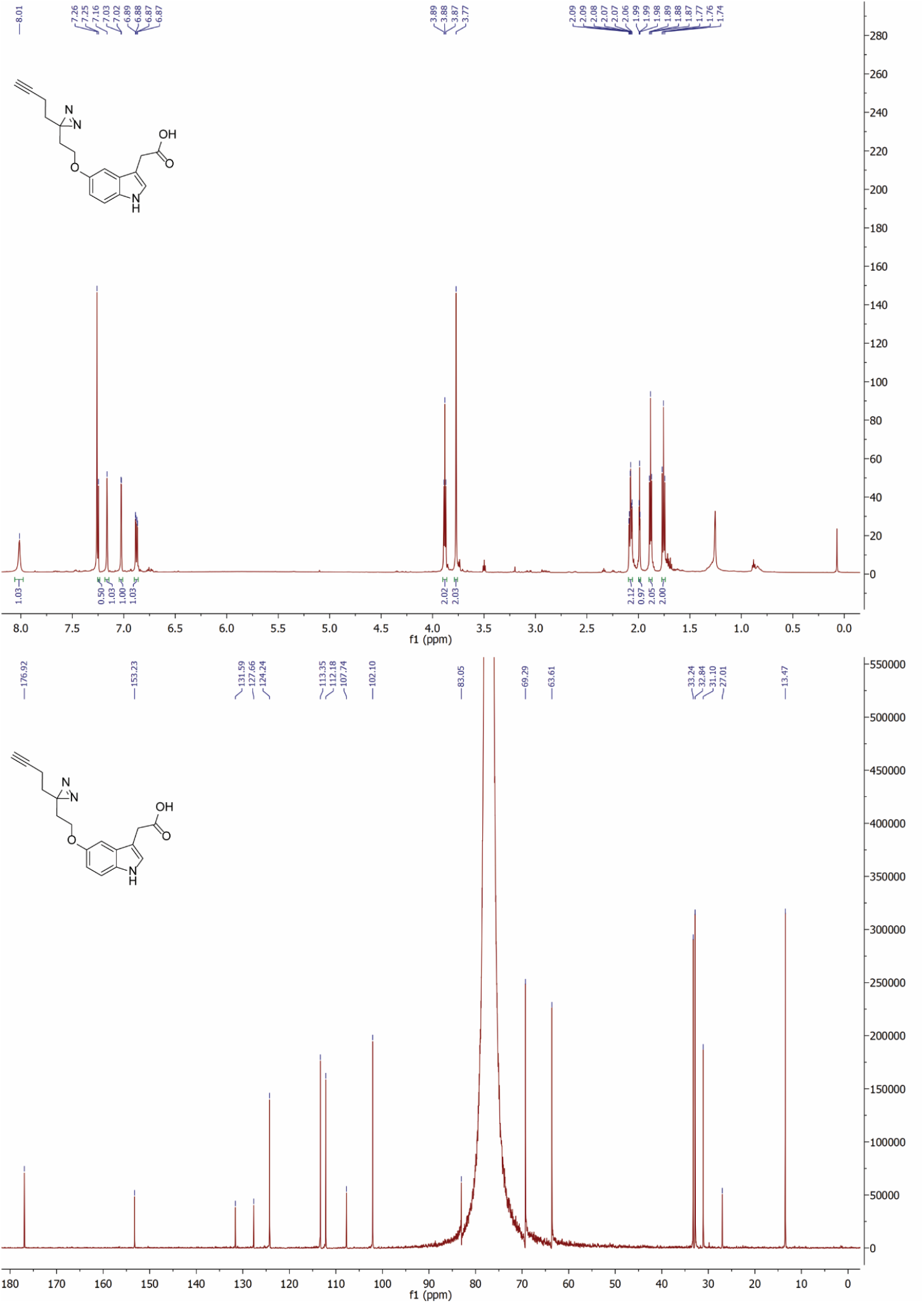

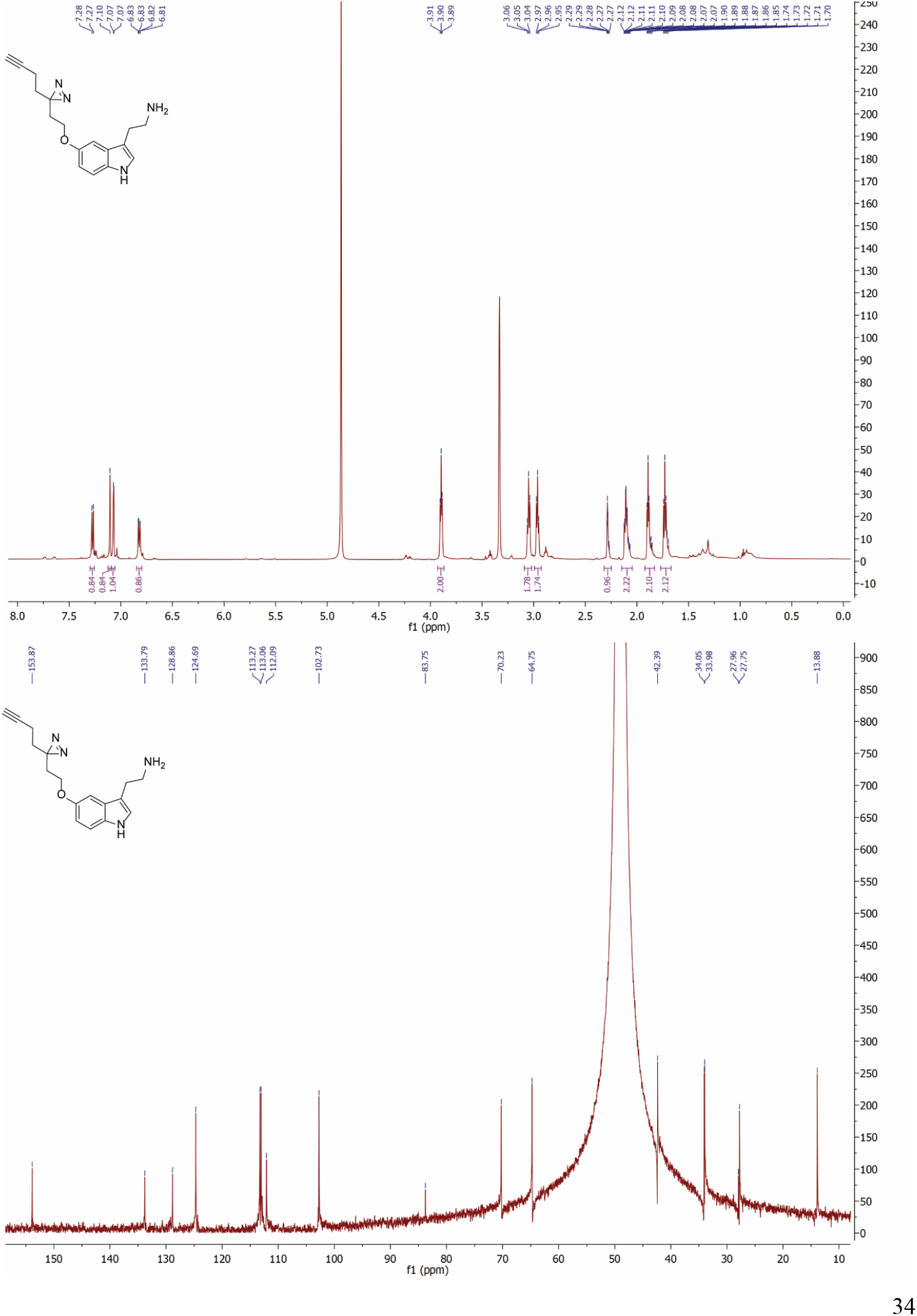

## MATERIALS AND METHODS

### Chemicals

Commonly used reagents were purchased from Sigma-Aldrich and Fisher Scientific. Specific chemicals were originated from following sources: 2,3,7,8-tetrachlorodibenzo-p-dioxin solution (TCDD, Sigma-Aldrich, Catalog #48599), indole-3-acetatic acid (Fisher Scientific, Catalog #11453194), tryptamine (Sigma-Aldrich, Catalog #76706), 2-phenethylamine (Sigma-Aldrich, Catalog #42346), tyramine (Sigma-Aldrich, Catalog #80345), histamine (Sigma-Aldrich, Catalog #59964), 5-fluorotryptamine hydrochloride (AstaTech, Catalog #52030), 6-fluorotryptamine (AstaTech, Catalog #W10003), 7-fluorotryptamine hydrochloride (Cayman Chemical, Catalog #17323), 7-chlorotryptamine hydrochloride (Santa Cruz Biotechnology, Catalog #sc-268384), 7-bromotryptamine hydrochloride (Santa Cruz Biotechnology, Catalog #sc-268370), *N*-methyltryptamine (Cayman Chemical, Catalog #12067), (1H-Indol-3-yl)methanamine (Sigma-Aldrich, Catalog # CDS020852), α-methyltryptamine hydrochloride (Cayman Chemical, Catalog #15692), α-ethyltryptamine (Cayman Chemical, Catalog #15671), L(-)-epinephrine (Fisher Scientific, Catalog #AC204400010), clonidine hydrochloride (Tocris, Catalog #0690), pyridoxal-5’-phosphate (Oakwood Chemical, Catalog #044568), azido-PEG3-biotin (Click Chemistry Tools, Catalog #AZ104).

The synthesis of alk-IAA, x-alk-IAA and x-alk-TA was according to reported synthetic procedures.^63,64^ The product was purified by silica gel flash column chromatography, determined by NMR (Bruker 600 MHz) and LC-HRMS (Orbi XL). alk-IAA: ^1^H-NMR (600 MHz, CDCl3) δ (ppm) 10.00 (s, 1H), 7.31-7.30 (d, 1H), 7.28 (s, 1H), 7.23-7.22 (d, 1H), 6.85-6.83 (dd, 1H), 4.76-4.75 (d, 2H), 3.71 (s, 2H), 2.99-2.98 (t, 1H); ^13^C-NMR (150 MHz, CDCl3) δ (ppm) 173.14, 152.84, 133.19, 128.88, 125.51, 112.98, 112.75, 108.92, 103.82, 80.56, 76.30, 57.30, 31.39. HRMS (ESI) calculated for C13H11NO3 [M+H]^+^: 230.0817, found 230.0816.

x-alk-IAA: ^1^H-NMR (600 MHz, CDCl3) δ (ppm) 8.31 (s, 1H), 7.26-7.25 (d, 1H), 7.16 (s, 1H), 7.03-7.02 (d, 1H), 6.89-6.87 (dd, 1H), 3.89-3.87 (t, 2H), 3.77 (s, 2H), 2.09-2.06 (m, 2H), 1.99-1.98 (t, 1H), 1.89-1.87 (t, 2H), 1.77-1.74 (t, 2H); ^13^C-NMR (150 MHz, CDCl3) δ (ppm) 176.92, 153.23, 131.59, 127.66, 124.24, 113.35, 112.18, 107.74, 102.10, 83.05, 69.29, 63.61, 33.24, 32.84, 31.10, 27.01, 13.47. HRMS (ESI) calculated for C17H17N3O3 [M+H]^+^: 312.1348, found 312.1341.

x-alk-TA: ^1^H-NMR (600 MHz, CDCl3) δ (ppm) 7.28-7.27 (d, 1H), 7.10 (s, 1H), 7.07 (s, 1H), 6.83-6.81 (dd, 1H), 3.91-3.89 (t, 2H), 3.06-3.04 (t, 2H), 2.97-2.95 (t, 2H), 2.29-2.27 (m, 1H), 2.12-2.07 (m, 2H), 1.90-1.85 (m, 2H), 1.74-1.70 (m, 2H); ^13^C-NMR (150 MHz, CDCl3) δ (ppm) 153.87, 133.79, 128.86, 124.69, 113.27, 113.06, 112.09, 102.73, 83.75, 70.23, 64.75, 42.39, 34.05, 33.98, 27.96, 27.75, 13.88. HRMS (ESI) calculated for C17H20N4O [M+H]^+^: 297.1715, found 297.1717.

### Cell Culture

HEK293T and HT-29 cells were obtained from ATCC. HTLA cells (a HEK293 cell line stably expressing a tTA-dependent luciferase reporter and a β-arrestin2-TEV fusion gene) were a gift from the laboratory of Gilad Barnea at Brown University. HEK293T and HT-29 cells were cultured in DMEM (Gibco) supplemented with 10% FBS (GE Healthcare HyClone) and 100 U/mL penicillin-streptomycin (ThermoFisher). HTLA cells were cultured in DMEM supplemented with 10% FBS and 100 U/mL penicillin-streptomycin, 2 μg/mL puromycin and 100 μg/mL hygromycin B. All cells were cultured in a 37 °C incubator with 5% CO_2_.

### Luciferase reporter assay

HEK293T cells were seeded in CoStar white 96-well plate (Corning) at 20,000 cells/well for 16 h. The cells were transiently transfected with three plasmids including pCMV3-N-HA-AhR (Sinobiological, HG10456-NY), pGL4.43[luc2P/XRE/Hygro] (Promega, E4121) and pRL-TK (Promega, E2241) using Lipofectamine 3000 Transfection Reagent (Invitrogen, L3000-015). After 24 hours, cells were stimulated with 0.1% DMSO, 10 nM TCDD, 100 μM IAA, alk-IAA or x-alk-IAA for 16 hours. Luminescence was detected after addition of dual-luciferase reporter assay reagent (Promega, E1960), using a BioTek Synergy Neo2 Multi-Mode Microplate Reader with a 0.5 second integration time. Firefly luciferase luminescence normalized to the Renilla luciferase control is shown, with error bars indicating the S.E.M. for three replicates.

### In-gel fluorescence and Western blot

For in-gel fluorescent labeling, HEK293T cells were grown in 6-well plate at 500,000 cells/well to above 90% confluence before transfection with respective plasmids pCMV3-N-HA-AhR, pcDNA3.1_ADRA2A-Tango (Addgene 66216), pcDNA3.1_GPRC5A-Tango (Addgene 66382), pcDNA3_GPRC5A-Flag, pcDNA3_GPRC5A_N252A-Flag, pcDNA3_GPRC5A_F256A-Flag, or pcDNA3_GPRC5A_N252A_F256A-Flag. After 24 h, cells were change to OMEM medium containing 0.1% DMSO, 100 μM x-alk-IAA, or 100 μM x-alk-TA, and incubated for 1 h at 37 °C. Then the cells were washed and incubated in cold DPBS on ice, exposed to 365 nm UV light for 10 min. Cells were lysed in 500 μL of lysis buffer including 20 mM HEPES buffer, pH 7.5, 150 mM NaCl, 1% NP-40, 1x cOmplete protease inhibitor cocktail (Sigma-Aldrich) for 15 min. Cell lysates were cleared by centrifugation at 20,000g for 20 min, and the soluble proteins were incubated with EZview red anti-HA or anti-Flag affinity gel (Sigma) for 3 h at 4 °C. The immunoprecipitated proteins were labeled with azido-rhodamine via click chemistry, resolved by SDS-PAGE, and visualized on a Bio-Rad ChemiDoc MP imaging system. For in-gel fluorescent profiling, HT-29 cells were cultured in 6-well plate at 500,000 cells/well to above 90% confluence, treated with 0.1% DMSO, 100 μM alk-IAA, 100 μM x-alk-IAA, or 100 μM x-alk-TA with or without UV irradiation. Cells were lysed in 1% SDS lysis buffer, labelled by azido-rhodamine, resolved by SDS-PAGE and visualized. For Western blot analysis, cell lysates prepared as above were labeled with azido-PEG3-biotin (Click Chemistry Tools, AZ104), incubated with high-capacity neutravidin beads (Thermo Scientific Pierce), resolved by SDS-PAGE and analyzed by Western blot. Antibodies for ADRA2A (14266-1-AP), GPRC5A (10309-1-AP), GPR107 (25076-1-AP), GPR108 (24009-1-AP), STING/TMEM173 (19851-1-AP), MAVS (14341-1-AP), SLC16A3/MCT4 (22787-1-AP) were purchased from Proteintech, SCL18B1 (bs-9530) and SLC38A2 (bs-12125R) from Bioss.

### Chemical proteomics

HT-29 cells were grown in 10-cm dish to above 90% confluence, treated with DMSO, alk-IAA or x-alk-IAA with or without UV-irradiation, lysed and labeled with azido-PEG3-biotin as shown above. The proteins enriched by neutravidin beads were digested with MS-grade trypsin (Promega). The on-beads digested peptides were collected and desalted by the in-house prepared reversed phase C18 Stage-tip, dried by Speed-vac, redissolved in 0.1% TFA/H_2_O. Tryptic peptides form each of the 4 replicates were separated by the reversed phase C18 column on an EASY-nLC 1000 (Thermo Scientific) system at a flow rate of 200 nL/min using a 70 min gradient from 5% B to 45% B (A: 0.1% formic acid, B: acetonitrile/0.1% formic acid) followed by a washing step at 100% B. Mass spectra were collected on an LTQ-Orbitrap Fusion Tribrid mass spectrometer (Thermo Scientific). Full scan MS spectra were acquired in the Orbitrap at a resolution of 60,000 (at m/z 300). The raw data were analyzed using MaxQuant (version 1.6.0.16)^65^. Parameters were used as default settings for label-free quantification (LFQ). For protein identification, the false discovery rate was set to 0.01, the minimum peptide length allowed was seven amino acids, and the minimum number of unique peptides allowed was set to one. Further data processing was performed using Perseus (version 1.6.0.7).^66^ LFQ intensities were log2 transformed. The data were filtered to remove reverse contaminants, only identified by sites. Proteins were filtered to be detected in at least 3 valid values out of 8 experiments. The replicates were grouped together, and empty values were imputed with random numbers from a normal distribution (width = 0.3 and shift = 1.8). Significantly enriched proteins were identified as having a p-value below 0.05 and at least two folds change when comparing LFQ intensities of chemical reporter-treated group with DMSO vehicle group. The data was visualized in a volcano plot of log2 ratio change versus the negative logarithmic p values. Gene ontology analysis of the protein hits was performed by Enrichr.

### PRESTO-Tango assay

The PRESTO-Tango plasmids were a gift from Bryan Roth (Addgene 1000000068).^43^ HTLA cells were seeded in a CoStar white 96-well plate (Corning) at 20,000 cells/well for 16 h. The cells were transiently transfected with individual GPCR-Tango plasmid using Lipofectamine 3000 Transfection Reagent (Invitrogen, L3000-015). After 24 h, cells were stimulated with various molecules of different concentration for 16 h. Cell culture medium were changed with 50 μL Bright-Glo luciferase assay system reagent (Promega) that was firstly diluted 20 folds with 1x DPBS buffer. After incubation for 20 min at room temperature in dark, luminescence was detected using a BioTek Synergy Neo2 Multi-Mode Microplate Reader with a 0.5 second integration time.

### Bacterial culture

Some commercially available bacteria species and strains are sourced as follows: *E. coli* DH5a (NEB), *E. faecalis* OG1RF (ATCC 47077), *E. faecalis* PCI 1325 (ATCC 14506), *E. faecalis* V583 (ATCC 700802), *E. faecium* DO/TX0016 (ATCC BAA-472), *E. faecium* (ATCC 700221), *E. faecium* TX0082 (BEI Resources), *E. faecium* ERV165 (BEI Resources), *E. durans* 23C2 (ATCC 6056), *E. hirae* R (ATCC 8043), *E. mundtii* NCDO 2375 (ATCC 43186), *E. gallinarum* NCDO 2313 (ATCC 51559), *M. morganii* ATCC 25830 (Microbiologics), *S. capitis* ATCC 35661 (Microbiologics), *S. epidermidis* ATCC 14990 (Microbiologics), *L. brevis* ATCC 14869 (Microbiologics). *E. faecium* Com15 was provided as a gift by Michael Gilmore (Harvard Medical School), *R. gnavus* ATCC 29149 by Sean F. Brady (The Rockefeller University), *C. sporogenes* ATCC 15579 by Chun-Jun Guo (Weill Cornell Medical School). Bacteria were cultured in the anaerobic chamber (Coy Laboratory Products) at 37 °C for 48 h or bacterial shaker incubator of 220 rpm at 37 °C for 16 h in LB, BHI, TSB or MRS media respectively. Bacterial growth was determined by optical density measurement. Bacterial cultures were pelleted by centrifugation at 5,000 g, supernatants were filtered through 0.2 μM membrane (Millipore) followed by 3,000 Da MWKO membrane (Amicon).

### Phylogenetic analysis

The analysis of TyrDC orthologs across different species in the human microbiome was conducted as previously described.^9^ Briefly, GenBank coding sequences from genome FASTA assemblies within the Human Microbiome Project^47^ were downloaded via BioProject Accession number PRJNA28331 and queried by BLASTP using TyrDC *E. faecium* Com15 (AZV35689.1). For genomes with multiple hits, the hit sequence with the highest percent identity with the query sequence was chosen as the TyrDC ortholog. Separately, GenBank genomic assemblies were queried for 16S rRNA sequences using Barrnap (GitHub.com/tseemann/barrnap). The longest predicted 16S rRNA sequence for each strain was used for alignment by Clustal Omega^67^ using default settings. The multiple sequence alignment was trimmed by TrimAl v1.3 (automated method)^68^, and an approximate maximum-likelihood phylogenetic tree was constructed using FastTree v2.1.10^69^. Data were visualized as a cladogram by iTOL.^70^ For the unrooted phylogenetic clustering, protein sequences were aligned by Clustal Omega with output in FASTA format. The FASTA alignment was then uploaded to the IQ-TREE tool using default settings with the auto substitution model, and 10,000 bootstrap alignments were calculated using the ultrafast setting^71^. The resulting phylogenetic tree file was visualized using iTOL.

### LC-MS analysis

Chemicals and membrane-filtered bacterial cultures were analyzed by 1290 Infinity II LC/MSD system (Agilent technologies) using Poroshell 120 EC-C18 column (2.1 x 50 mm, 4 μM). The flow rate was 0.5 mL/min using 0.1% formic acid in water as mobile phase A and 0.1% formic acid in acetonitrile as mobile phase B at room temperature. The gradient was applied as below: 0-2 min: 5% B, 2-5 min: 5-95% B, 5-7 min: 95% B, 7-10 min: 95-5% B. MSD API-ES SIM mode on sample target masses of tryptamine (m/z = 161.1), tyramine (m/z = 138.1), phenethylamine (m/z = 122.1), histamine (m/z 112.1). Extracted intensity and retention time were analyzed by GraphPad Prism 8.

### Generation of *TyrDC-KO E. faecium* Com15 strains

*TyrDC* knockout of *E. faecium* Com15 was generated using CRISPR-Cas9 mediated recombineering as previously described.^48^ In brief, pRecT containing *E. feacium* Com15 was co-electroporated with ssDNA (Integrated DNA technologies) encoding homology arms targeting *TyrDC*, and pCas9 harboring gRNA targeting the *TyrDC* deletion region. The resulting knockout strain carries a 74 base-pair deletion in the *TyrDC* gene, verified via PCR and sanger sequencing (Genewiz). The strain was then cured of pRecT and pCas9 through passaging to generate isogenic mutants. Single colony was inoculated for preparation of glycerol stock. pKH12-*TyrDC* was generated by cloning his-tagged *E. faecium TyrDC* into psrtA1 with *srtA-1* omitted from the plasmid. The resulting plasmid contains a *TyrDC* under the constitutive *bacA* promoter in a pKH12 backbone^72^. pKH12-*TyrDC* or vector pKH-12 was transformed into *E. feacium* Com15 through electroporation. Single colony was inoculated and cultured in BHI media for 16 h. Cleared bacterial cultures were filtered, analyzed by LC-MS and PRESTO-Tango assay.

### Cloning and analysis of aAAs decarboxylase

*TyrDC* was amplified from *E. faecium* Com15 genomic DNA, *TrpDC* from *R. gnavus* ATCC 29149, *Gln/TyrDC* from *M. morganii* ATCC 25830, and inserted to pKH12 vector by Gibson assembly (NEB), respectively. The plasmids were transformed into chemical competent *E. coli* DH5α (NEB). Single colony was inoculated in 5 mL of LB medium containing 25 μg/mL chloramphenicol for 16 h, and was then transferred to into Minimal media supplemented with 10 mM Trp, 2 mM Tyr, 10 mM Phe or 10 mM His of a OD_600_ value of 0.1. After 16 h, bacteria were pelleted, the supernatants were filtered through 0.2 μM membrane followed by 3,000 Da MWKO membrane. The filtered cultures were analyzed by LC-MS and PRESTO-Tango assay.

### Analysis of *E. faecium* Com15 TyrDC enzymatic activity

For preparation of TyrDC, *E. coli* DH5α transformed with pKH12-*TyrDC* were grown in 500 mL LB medium containing 25 μg/mL chloramphenicol starting from OD_600_ of 0.1 for 24 h at 37 °C with shaking at 220 rpm. Cells were pelleted, resuspended in 40 mL 50 mM Tris pH 8 containing 0.25 M NaCl, lysed by sonication. The lysate was clarified by centrifugation, incubated with 2 mL of Ni-NTA resin at 4 °C for 16 h. The resins were loaded on a column, washed three times with lysis buffer, eluted using a gradient of 25 mM to 200 mM imidazole in lysis buffer. Fractions containing pure protein were combined and dialyzed over two rounds into in 50 mM Tris pH 8 containing 0.20 M NaCl and 10% w/v glycerol. The dialyzed protein was concentrated using spin columns, concentration was determined by BSA assay, protein aliquots were frozen in liquid nitrogen and stored at −80 °C.

Purified TyrDC was incubated with pyridoxal-5’-phosphate (PLP) in the reaction buffer (200 mM sodium acetate buffer, pH 5.5) for five minutes. The enzymatic reaction was initiated by adding the preincubated TyrDC-PLP mix with substrates dissolved in the reaction buffer to yield the final concentrations for 100 μM PLP, 100 nM enzyme and 10 μM-2000 μM tyrosine or 1 mM-60 mM phenylalanine at room temperature for 1 min or 37 °C for 10 min, respectively. Aliquots of the reaction were quenched by diluting 10-fold into ice cold methanol, centrifugated and 2 μl of the supernatant was analyzed. Rates were calculated and fit to a standard Michaelis–Menten kinetics curve in Graphpad prism 8.

